# Long-chain fatty acyl-coenzyme A activates the mitochondrial fission factors MiD49 and MiD51 by inducing their oligomerization

**DOI:** 10.1101/2023.07.31.551267

**Authors:** Ao Liu, Frieda Kage, Gracie Sapp, Halil Aydin, Henry N. Higgs

## Abstract

Mitochondrial fission occurs in many cellular processes, but the regulation of fission is poorly understood. We show that long-chain acyl coenzyme A (LCACA) activates two related mitochondrial fission proteins, MiD49 and MiD51, by inducing their oligomerization, activating their ability to stimulate DRP1 GTPase activity. The 1:1 stoichiometry of LCACA:MiD in the oligomer suggests interaction in the previously identified nucleotide-binding pocket, and a point mutation in this pocket reduces LCACA binding and LCACA-induced oligomerization for MiD51. In cells, this LCACA binding mutant does not assemble into puncta on mitochondria or rescue MiD49/51 knock-down effects on mitochondrial length and DRP1 recruitment. Furthermore, cellular treatment with the fatty acid analogue 2-bromopalmitate, which causes increased acyl-CoA, promotes mitochondrial fission in an MiD49/51-dependent manner. These results suggest that LCACA is an endogenous ligand for MiDs, inducing mitochondrial fission and providing a potential mechanism for fatty acid-induced mitochondrial fragmentation. Finally, MiD49 or MiD51 oligomers synergize with MFF, but not with actin filaments, in DRP1 activation, suggesting distinct pathways for DRP1 activation.

## Introduction

Mitochondrial fission is a key cellular process, with defects in fission being linked to multiple human diseases^1^. The reasons for which mitochondria undergo fission vary considerably, including for distribution to daughter cells during cell division, distribution to remote areas of polarized cells, and in response to changing metabolic conditions^2,3^. In addition, mitochondrial fission is an important step in mitophagy of damaged mitochondrial segments, and defects in fission result in decreased overall mitochondrial health^4^. Finally, mitochondrial fission proteins participate in the production of mitochondrially-derived vesicles^5^, an additional mechanism for removal of dysfunctional mitochondrial components.

A key protein in mitochondrial fission is the membrane-remodeling GTPase DRP1, which is recruited from the cytoplasm to the outer mitochondrial membrane (OMM), where it oligomerizes into a ring structure around the mitochondrion^2^. Oligomerization activates DRP1’s GTPase activity by bringing the GTPase domains in close proximity^6–8^. GTP hydrolysis by the DRP1 oligomer results in constriction of the oligomer, driving the fission process.

Given the variety of cellular situations requiring mitochondrial fission, it is likely that there are multiple mechanisms for regulating the process. One potential step for differential control of mitochondrial fission is in DRP1 recruitment to the OMM. In mammals, three DRP1 receptors have been identified: mitochondrial fission factor (MFF), and two related proteins: mitochondrial dynamics proteins of 49 and 51 kDa (MiD49 and MiD51, also called MIEF2 and MIEF1 respectively)^2^. All three proteins are widely expressed in mammalian cells. However, it is unclear to what extent these receptors function together versus independently. Knock-down or knock-out of MFF alone causes dramatic mitochondrial elongation in many metazoan cell types tested^9–13^ with one exception^14^, suggesting that it might play a role in many forms of mitochondrial fission. Knock-down or knock-out of MiD49 and MiD51 also causes mitochondrial elongation, although the effect is more variable between studies and there are differing reports of the redundancy between MiD49 and 51^11,13–16^.

These findings suggest that MiD49 and MiD51 might play roles in a sub-set of mitochondrial fission events. Three other properties of MiD49/51 suggest that they act in a context-specific manner. First, while depletion of MiD49/51 results in mitochondrial elongation, over-expression of either protein has the same effect ^11,15–17^. Over-expressed MiD49/51 also causes extensive DRP1 recruitment to mitochondria ^11,15–17^, suggesting that unregulated MiD-mediated DRP1 recruitment is detrimental to its controlled assembly during fission. Second, MiD49/51 expressed at low levels appear punctate on mitochondria, suggesting oligomerization ^13,17,18^. In contrast, the purified cytoplasmic regions of either MiD49 or MiD51 behave as monomers or dimers^19,20^, suggesting that this oligomerization is regulated.

A third property of both MiD49 and MiD51 is that their purified cytoplasmic regions do not stimulate DRP1 GTPase activity biochemically^19,21^, in contrast to MFF^22–24^. These findings suggest that the MiD proteins themselves require activation in order to activate DRP1. The structures of both proteins reveal large putative ligand-binding cavities (**Fig. 1a**). MiD51 is capable of binding purine nucleotide diphosphate, with ADP being preferred to GDP^19,20^, while MiD49 displays no affinity for these ligands^20^. The effect of ADP on MiD51’s ability to activate DRP1 is modest, with an approximate 2-fold stimulation of DRP1 GTPase activity over DRP1 alone ^20^. These results suggest that MiD49 and MiD51 might bind alternate ligands to stimulate their DRP1-activating ability in cells.

**Fig. 1:**
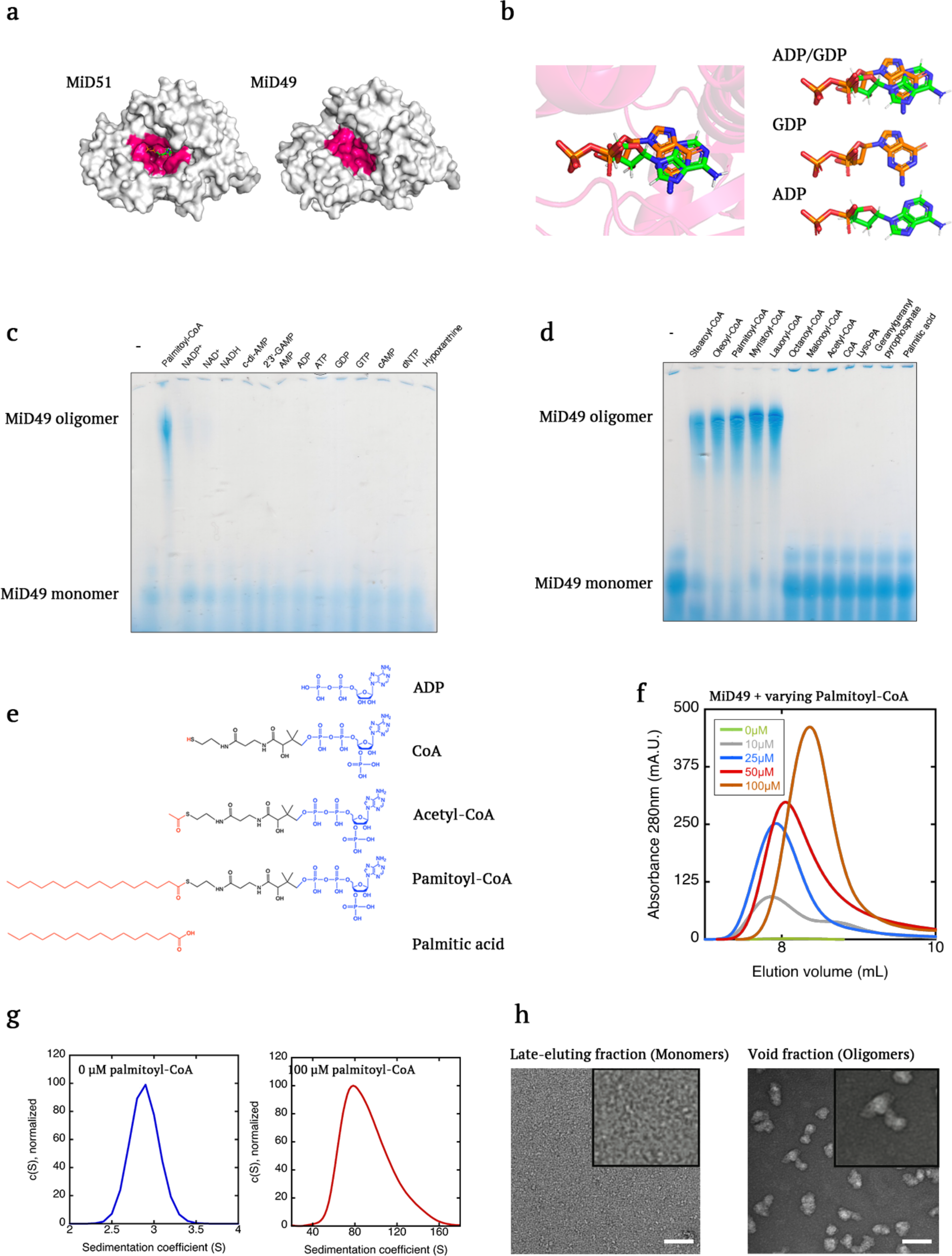
Long-chain acyl CoA induces MiD49 oligomerization. **a,** Surface model of human MiD51 with bound ADP (PDB 4NXW, left) and murine MiD49 without bound ligand (PDB 4WOY, right). Residues lining the binding pocket in pink. **b,** Configurations of ADP and GDP bound to human MiD51, from PDB 4NXW and 4NXX. **c,** Blue-native gel electrophoresis of MiD49 cytoplasmic region (100 µM) mixed with 500 µM of the indicated purine-containing compounds. **d,** Blue-native gel electrophoresis of MiD49 cytoplasmic region (100 µM) mixed with 500 µM of the indicated molecule. **e,** Bond-line formulas comparing ADP, CoA, acetyl-CoA, palmitoyl-CoA and palmitate. **f,** Size exclusion chromatography of MiD49 (100 µM) incubated with the indicated concentrations of palmitoyl-CoA. Peak shown is near the void volume of the Superose 6 column, indicative of a high molecular weight species. Full chromatogram in Extended Data Fig. 1c. **g,** Velocity analytical ultracentrifugation of peak fractions from Superose 6 size exclusion chromatography of MiD49 cytoplasmic region (100 µM) incubated without or with palmitoyl-CoA (100 µM). Sedimentation coefficients shown (2.9 and 82 S for 0 and 100 µM palmitoyl-CoA, respectively). C(S) normalized to the maximum peak value. Extended Data Fig. 1d shows molecular mass conversion. **h,** Negative stain electron microscopy of the void fraction from the 50 µM palmitoyl-CoA condition or the late-eluting fraction from the 0 µM palmitoyl-CoA condition from panel F. Scale bar, 150 nm.

We have previously shown that actin filaments can bind and activate DRP1^25,26^, and that inhibition of actin polymerization or of the actin polymerization factor INF2 decreases mitochondrial DRP1 recruitment and inhibits mitochondrial fission in cells^25,27,28^. Actin activates DRP1 GTPase activity in a characteristic ‘bi-phasic’ manner, in which low concentrations stimulate while higher concentrations do not^24,26^. This bi-phasic behavior is likely due to the fact that DRP1 activation requires inter-molecular binding between GTPase domains^6–8^, with low actin concentrations facilitating this juxtaposition while higher actin concentrations cause DRP1 to bind more sparsely along the filament, minimizing interaction. In addition to its ability to increase DRP1 activity in a stand-alone manner, actin can synergize with MFF in DRP1 activation, decreasing the concentration of MFF needed for DRP1 stimulation^24^. It is not clear what effect MiD49 or MiD51 would have on actin- or MFF-mediated DRP1 activation.

While mitochondrial fission is often associated with mitochondrial dysfunction and mitophagy, fission has also been correlated with increased mitochondrial fatty acid oxidation in several contexts, including brown adipocytes, hepatocytes, and pancreatic beta cells^3,29^. An obligatory step in mitochondrial long-chain fatty acid import is coupling of fatty acid to coenzyme A to make long-chain acyl-coenzyme A (LCACA). In this paper, we show that both MiD49 and MiD51 bind LCACA, which induces MiD oligomerization. LCACA-oligomerized MiD49 or MiD51 activates DRP1 GTPase activity ∼10-fold, and this activation is synergistic with MFF-mediated DRP1 activation but not with actin-mediated activation. An MiD51 mutant defective in LCACA binding does not assemble into punctate structures on mitochondria, recruit DRP1 to mitochondria, or rescue the mitochondrial elongation phenotype caused by MiD49/51 knock-down. Overall, our work suggests that MiD49 and MiD51 might respond to LCACA for DRP1 recruitment *en route* to mitochondrial fission.

## Results

### Long-chain acyl-CoA induces MiD49 oligomerization

Both MiD49 and MiD51 contain putative nucleotide-binding pockets structurally similar to that of cyclic GMP-AMP synthase (cGAS) (**Fig. 1a**). While MiD51 can bind ADP or GDP in this pocket, MiD49 displays no apparent binding to these purine nucleotides^20^. Interestingly, the purine rings of ADP and GDP adopt distinct orientations in the binding pocket of MiD51 (**Fig. 1b**).

The absence of a strong effect of ADP or GDP on MiD51’s ability to stimulate DRP1^19,20^, coupled with the absence of a known ligand for MiD49, prompted us to screen for other possible ligands. We postulated that a ligand for MiD49 might induce oligomerization, because of the punctate appearance of MiD49 in cells. Using blue-native gel electrophoresis (BNGE) on a murine MiD49 cytoplasmic region construct (amino acids 125-454, **Extended Data Fig. 1a, b**), we screened several purine-containing compounds for the ability to cause a shift in MiD49 mobility, indicative of higher oligomeric species. Of these compounds, only palmitoyl-coenzyme A (palmitoyl-CoA) causes such a mobility shift (**Fig. 1c**). Neither the fatty acid moiety alone nor CoA alone causes a similar shift (**Fig. 1d,e**). Long-chain acyl-CoAs (LCACA) including stearoyl (18 carbons), oleoyl (18 carbons, 1 double bond), palmitoyl (16 carbons), myristoyl (14 carbon) and lauroyl-CoA (12 carbon) cause a shift in MiD49 mobility (**Fig. 1d**). In contrast, octanoyl-CoA (8 carbon) does not cause an MiD49 mobility shift, nor do two short-chain acyl-CoAs found in the cytoplasm, acetyl-CoA and malonoyl-CoA (**Fig. 1d**). We also tested lysophosphatidic acid (LPA), as well as geranylgeranyl-pyrophosphate (GG-PP), neither of which display an ability to shift MiD49 mobility (**Fig. 1d**).

To examine the oligomerization effect further, we used size exclusion chromatography (SEC). MiD49 cytoplasmic region alone (100 µM) elutes near the position of ovalbumin (∼ 45 kDa), suggestive of a monomer (**Extended Data Fig. 1c**). Palmitoyl-CoA concentrations from 10 to 100 µM cause a fraction of the protein to shift to a peak near the void volume, indicative of higher-order oligomers (**Fig. 1f, Extended Data Fig. 1c**). By velocity analytical ultracentrifugation, this void fraction sediments as a broad peak centered at 82 S (**Fig. 1g**), with a calculated mass of 4815 kDa (**Extended Data Fig. 1d**), which would suggest oligomers averaging 126 subunits. In contrast, the late-eluting fraction sediments at 2.9 S (**Fig. 1g**), with an apparent mass of 36.4kDa (**Extended Data Fig. 1d**), which is close to the calculated monomer mass (38.7 kDa). Negative-stain transmission electron microscopy (EM) reveals that the void fraction contains a heterogeneous array of oblate particles, whereas the late-eluting fraction contains a more uniform spread of small particles (**Fig. 1h**). Analysis of a limited number of particles from the void fraction reveals long and short particle axes of 66.7±12.7 and 39.7±13.5 nm, respectively (**Extended Data Fig. 2a-c**), for a mean axial ratio of 0.61±0.20. For comparison, the majority of palmitoyl-CoA micelles are more spherical, with an axial ratio of 0.85±0.15 and a mean diameter of 9.5±1.8 nm (17 particles measured, **Extended Data Fig. 2d**).

**Fig. 2:**
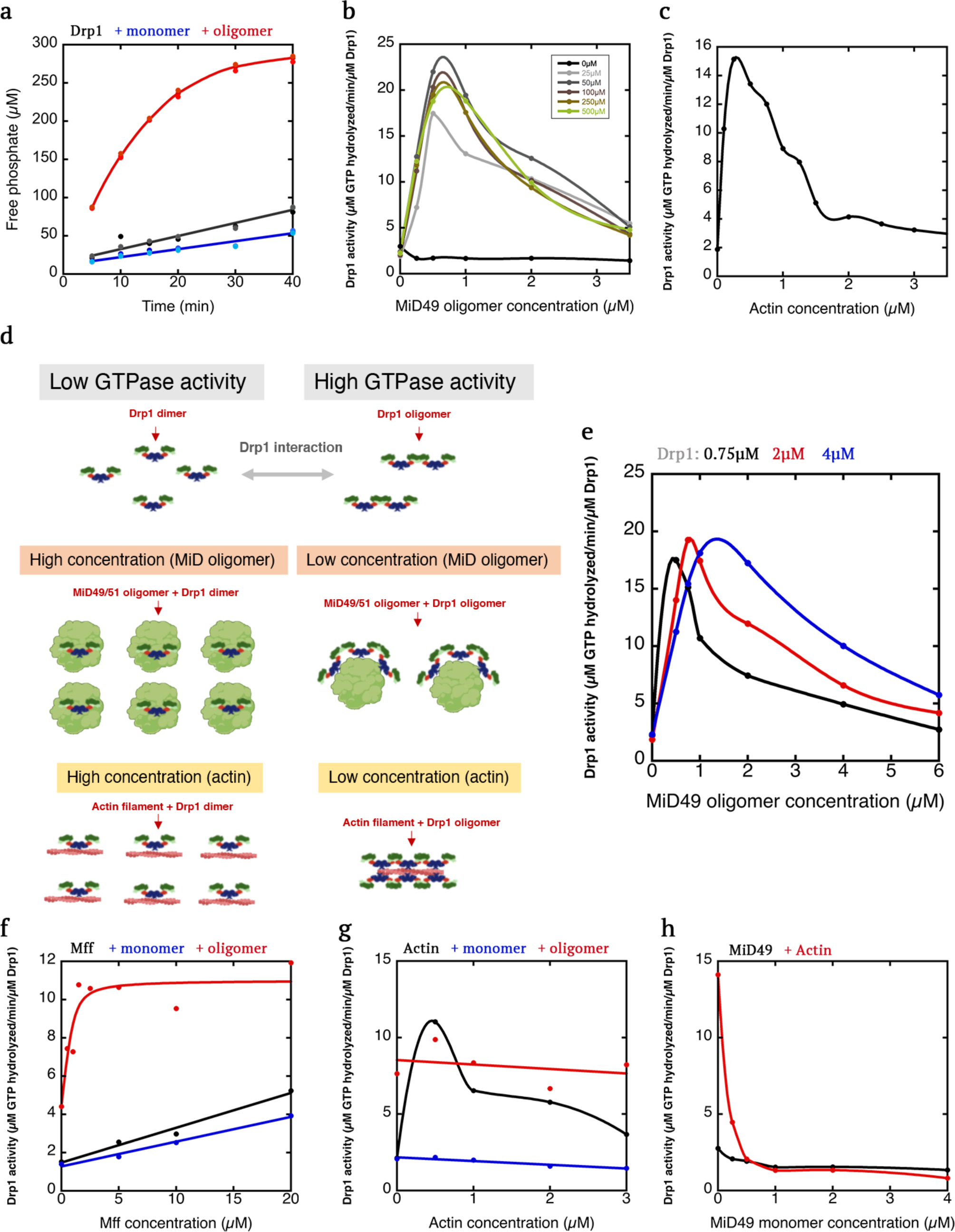
LCACA-induced MiD49 oligomers activate Drp1 in a synergistic manner with Mff. **a,** Drp1 GTPase assays (0.75 µM Drp1) alone (black points) or in the presence of 0.5 µM MiD49 oligomers (red) or monomers (blue). MiD49 oligomers from the 50 µM palmitoyl-CoA condition described in panel b. **b,** Effect of varying concentrations of MiD49 oligomers (assembled from mixtures of 100 µM MiD49 and varying concentrations of palmitoyl-CoA (given in µM)) on Drp1 GTPase activity (0.75 µM Drp1). Concentration effect of MiD49 monomers also shown. **c,** Effect of varying concentrations of actin filaments on Drp1 GTPase activity (0.75 µM Drp1). **d,** Schematic diagram of effect of MiD49 oligomers or actin filaments on Drp1 activity. At low concentration, MiD49 oligomers or actin filaments cause close juxtaposition of bound Drp1, allowing interaction between G domains and higher GTPase activity. At high concentration, Drp1 molecules are no longer juxtaposed, so that increased GTPase activity does not occur. **e,** Effect of varying concentrations of MiD49 oligomers (assembled from mixtures of 100 µM MiD49 and 50 µM palmitoyl-CoA) on Drp1 GTPase activity at 0.75, 2 and 4 µM Drp1. **f,** Effect of varying concentrations of Mff on Drp1 GTPase activity (0.75 µM Drp1) in the absence or presence of 250 nM MiD49 oligomers (red) or monomers (blue). Full curve to 100 µM Mff shown in Extended Data 5c. **g,** Effect of varying concentrations of actin filaments on Drp1 GTPase activity (0.75 µM Drp1) in the absence (black) or presence of 100 nM MiD49 oligomers (red) or monomers (blue). **h,** Effect of varying concentrations of MiD49 monomers on Drp1 GTPase activity (0.75 µM Drp1) in the absence (black) or presence (red) of 500 nM actin filaments.

The negative-stain EM also allows determination of the critical micelle concentration (cmc) for palmitoyl-CoA. From quantification of micelle number over a range of palmitoyl-CoA concentrations, we calculate a cmc of 47 µM in our buffer conditions (**Extended Data Fig. 2e**), similar to published values at similar ionic strength^30^. The fact that palmitoyl-CoA concentrations below this value cause a shift in MiD49 to by size-exclusion chromatography (**Fig. 1f**) suggests that MiD49 is not binding to palmitoyl-CoA micelles to affect this shift. In the ensuing experiments, we refer to the void peak as MiD49 oligomers, and the late-eluting peak as MiD49 monomers.

One question regards the stoichiometry of palmitoyl-CoA:MiD49 in the oligomer fraction. To quantify palmitoyl-CoA, we conducted reversed phase HPLC analysis. Palmitoyl-CoA elutes at 14.7 mL in this solvent system (**Extended Data Fig. 3a**), and the peak area (from 260 nm absorbance) is proportional to the quantity of palmitoyl-CoA loaded in the range of 0.5 – 2 nmole (**Extended Data Fig. 3b,c**). In an oligomer fraction containing 5.3 µM MiD49, the measured palmitoyl-CoA concentration is 5.5 µM (**Extended Data Fig. 3d**).

**Fig. 3.**
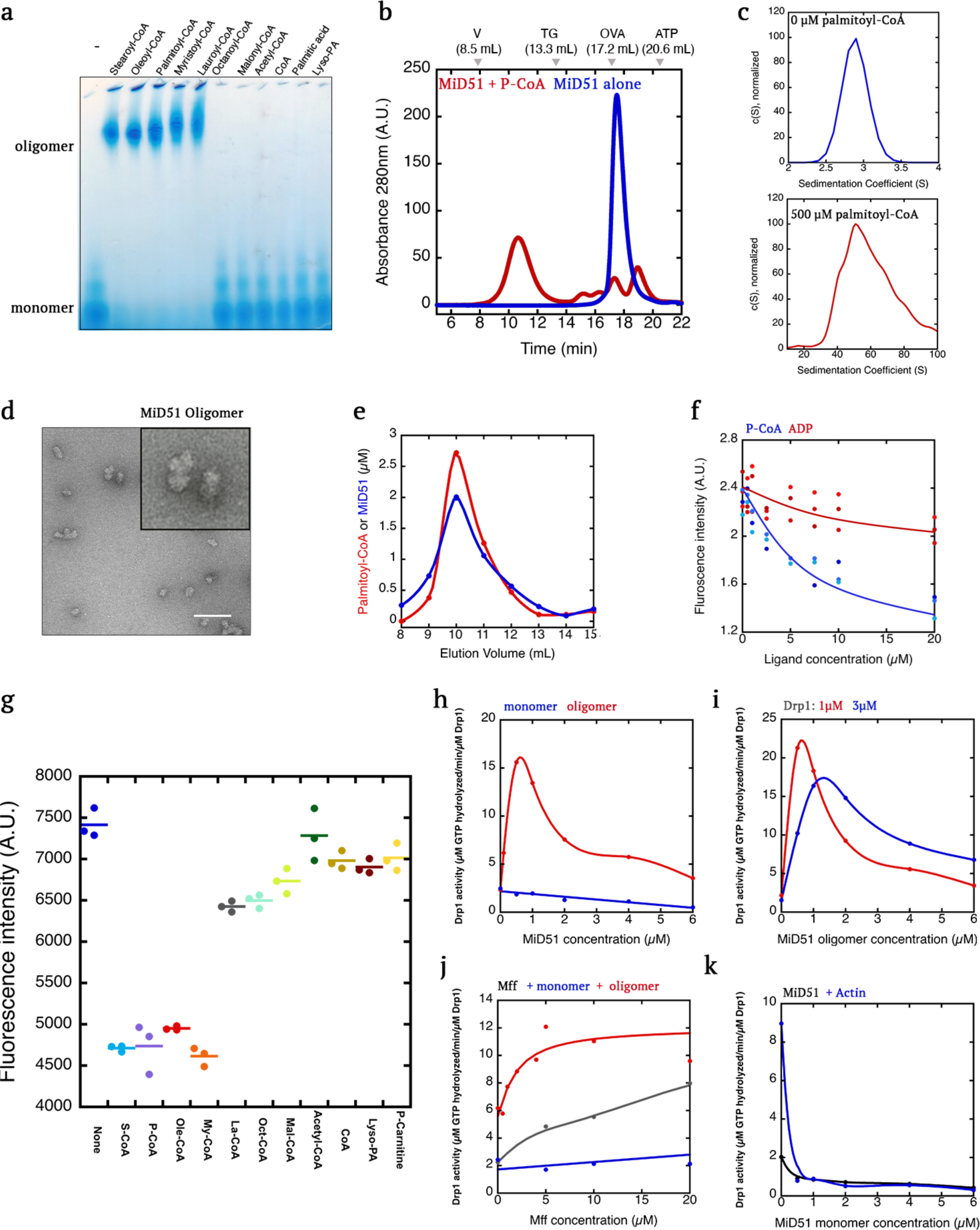
MiD51 oligomerizes in the presence of long-chain acyl-CoA. **a,** Blue-native gel electrophoresis of MiD51 cytoplasmic region (50 µM) alone or in the presence of 500 µM of the indicated molecules. **b,** Size-exclusion chromatography (Superose 6) of MiD51 (50 µM) alone or in the presence of 500 µM palmitoyl-CoA. **c,** Velocity analytical ultracentrifugation of the MiD51 oligomer peak from panel B (left) and the un-treated MiD51 monomer (right). MiD51 polypeptide concentration 5 µM in both cases. Calculated mass distribution shown in Extended Data Fig. 6c. **d,** Negative-stained electron micrograph of a representative particle from the MiD51 oligomer peak from panel B. More examples in Extended Data Fig. 2. **e,** Quantification of palmitoyl-CoA (phosphate assay) and MiD51 (Bradford assay) concentrations across the oligomer peak from panel b. **f,** MANT-ADP competition assay in which 300 nM MANT-ADP and 1 µM MiD51 are mixed with varying concentrations of palmitoyl-CoA (blue) or ADP (red) and the fluorescence intensity monitored. **g,** MANT-ADP competition assays in which 300 nM MANT-ADP and 1 µM MiD51 are mixed with 20 µM of the indicated molecule and the fluorescence intensity monitored. S-, P-, O-, M-, L-, Oct- and Mal-indicate stearoyl, palmitoyl, oleoyl, myristoyl, lauroyl-, octanoyl-, and malonyl-, respectively. **h,** Effect of varying concentrations of MiD51 oligomers on Drp1 GTPase activity (0.75 µM Drp1). Concentration effect of MiD51 monomers also shown. **i,** Similar experiment to that in Panel B, using two concentrations of Drp1, 1 µM and 3 µM. **j,** Effect of varying concentrations of Mff on Drp1 GTPase activity (0.75 µM Drp1) in the absence (gray) or presence of 100 nM MiD51 oligomers (red) or monomers (blue). **k,** Effect of varying concentrations of MiD51 monomers on Drp1 GTPase activity (0.75 µM Drp1) in the absence (black) or presence (blue) of 500 nM actin filaments.

As a second technique to determine palmitoyl-CoA concentration in MiD49 oligomer fractions, we used a phosphate assay (coenzyme A contains three phosphates). From these assays, the palmitoyl-CoA:MiD49 ratios from three oligomerization reactions range from 0.8 to 1.11 (**Extended Data Fig. 3e,f**). These results suggest that palmitoyl-CoA binds in a 1:1 complex with MiD49 in oligomers.

We also used the HPLC analysis method to test acyl chain preference of MiD49 further, by incubating MiD49 with equal concentrations of six acyl-CoAs (stearoyl, oleoyl, palmitoyl, myristoyl, lauroyl, and octanoyl), and then analyzing the acyl-CoA composition of the isolated MiD49 oligomer fraction. The three longer-chain acyl-CoAs (stearoyl, oleoyl, palmitoyl) are abundant in this fraction, while there is minimal myristoyl-CoA or lauroyl-CoA and no detectable octanoyl-CoA (**Extended Data Fig. 4**). This result suggests that, even though myristoyl-CoA or lauroyl-CoA can induce MiD49 oligomerization (**Fig. 1d**), they are out-competed by longer chain acyl-CoAs.

**Fig. 4.**
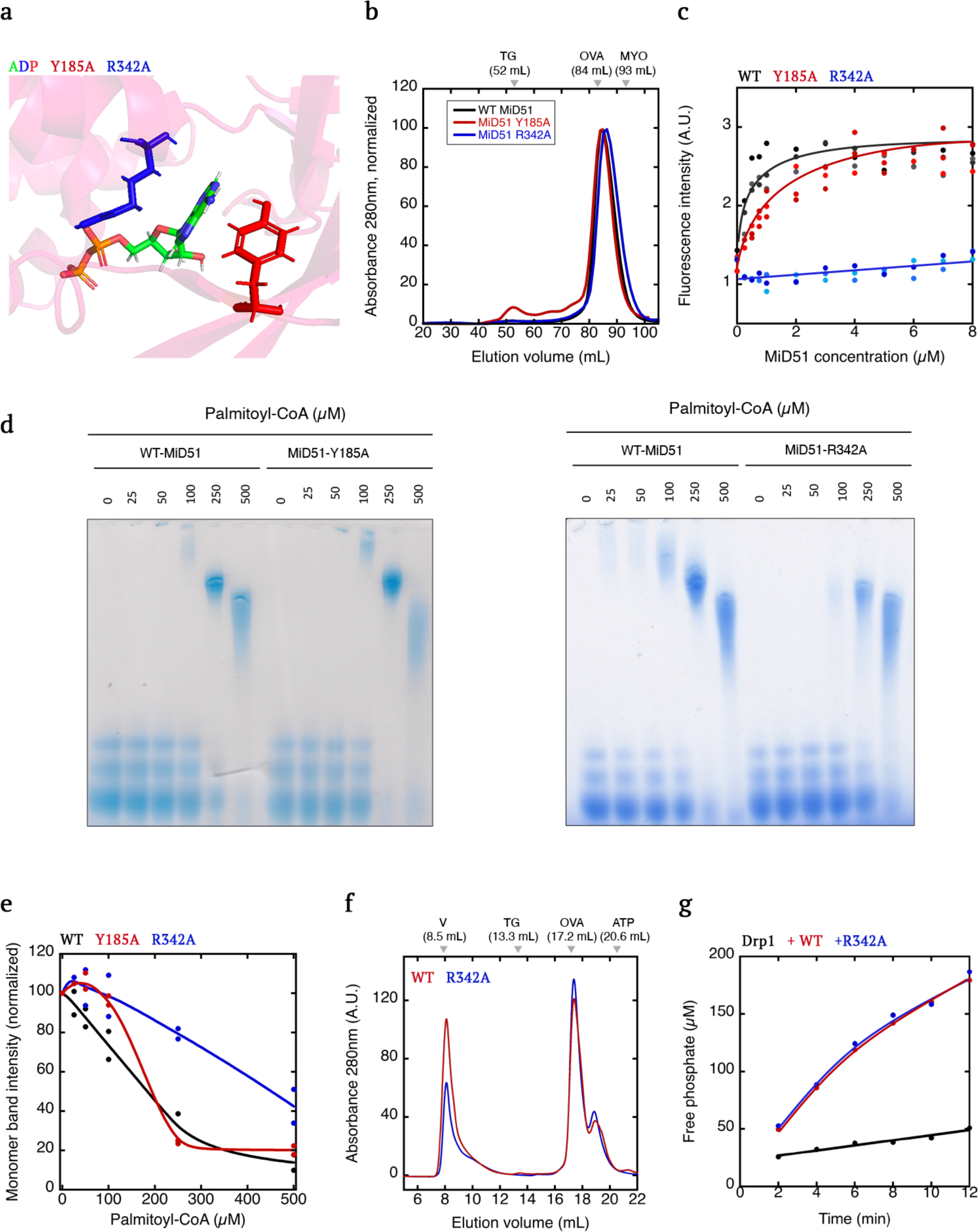
Mutation of R342 in MiD51 reduces LCACA-induced oligomerization. **a,** Model of human MiD51 with bound ADP (PDB 4NXW), showing the positions of R342 and Y185. **b,** Superdex 200 size exclusion chromatograms from the bacterial preparations of MiD51-WT, MiD51-R342A and MiD51-Y185A. Marker positions shown for: TG, thyroglobulin (660 kDa); OVA, ovalbumin (43 kDa); myoglobulin (17 kDa). **c,** MANT-ADP binding assay in which 300 nM MANT-ADP is mixed with the indicated concentrations of MiD51-WT, MiD51-R342A or MiD51-Y185A, and the fluorescence change monitored. **d,** Blue-native gel electrophoresis of MiD51-WT, MiD51-R342A or MiD51-Y185A (50 µM) mixed with varying concentrations of palmitoyl-CoA. **e,** Graph of density of the MiD51 monomer band as a function of palmitoyl-CoA concentration, from the blue-native gels such as in panel B (two independent gels for each construct). **f,** Superose 6 size exclusion chromatography of 50 µM MiD51-WT or MiD51-R342A mixed with 250 µM palmitoyl-CoA. Oligomer and monomer peaks indicated. **g,** Drp1 GTPase assays containing Drp1 alone (0.75 µM, black points) or in the presence of 2 µM MiD51-WT (red) or MiD51-R342A (blue).

### MiD49 oligomers stimulate DRP1 GTPase activity

The preceding results suggest that LCACA binds MiD49 in a 1:1 complex, and promotes MiD49 oligomerization. We next tested the effect of MiD49 oligomers on DRP1 GTPase activity. Similar to past results^19,20^, MiD49 monomers do not stimulate DRP1 GTPase activity (**Fig. 2a**). In contrast, MiD49 oligomers display an approximate 10-fold stimulation (**Fig. 2a**). We then analyzed the concentration dependence of DRP1 activation by MiD49 oligomers. Intriguingly, MiD49 oligomers have a bi-phasic effect on DRP1 GTPase activity, with stimulatory effects up to 1 µM and then decreasing activation at higher concentrations (**Fig. 2b**). This effect is similar for MiD49 oligomers isolated from a range of initial MiD49:palmitoyl-CoA ratios in the oligomerization reaction (**Fig. 2b**).

The bi-phasic effect of MiD49 oligomer concentration on DRP1 GTPase activity is reminiscent of that shown by actin filaments^26^ (**Fig. 2c, Extended Data Fig. 5a**). We postulate that, in both cases, low concentrations of the oligomeric molecule (MiD49 or actin filaments) induce the juxtaposition of DRP1 G domains, which stimulates GTPase activity^6–8^. Beyond a certain concentration of oligomeric molecule, however, DRP1 binding becomes sparser and G domains are separated (**Fig. 2d**). If this situation is true, increasing the DRP1 concentration should shift the optimally stimulating MiD49 oligomer concentration to a higher value. Indeed, the concentration of MiD49 oligomer needed to reach peak activation increases with increasing DRP1 concentration (**Fig. 2e**).

**Fig. 5.**
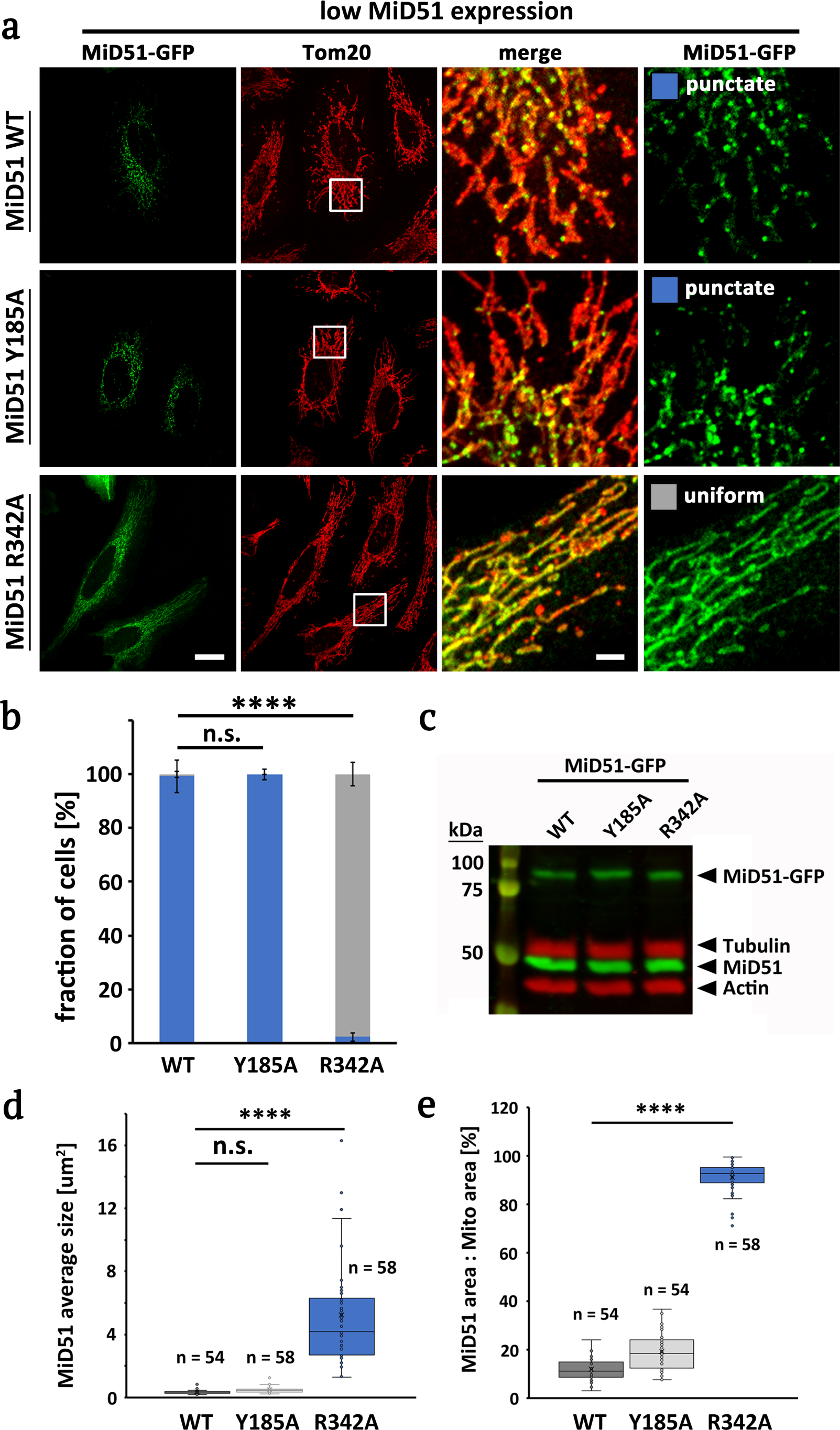
Effect of LCACA binding mutant on ability of MiD51 to form puncta in cells. **a,** GFP fusion constructs of MiD51-WT, MiD51-Y185A or MiD51-R342A were transiently transfected into HeLa cells, then cells were fixed and stained for mitochondria (Tom20). Low-expressing cells shown here (requiring 500 msec exposure at 100% laser power for GFP). Boxed regions denote insets shown in merges to the right of full images. Scale bars, 20 mm in full images and 3 mm in insets. **b,** Bar graph illustrating % cells displaying punctate (blue) or uniform (gray) mitochondrial GFP pattern for low-expressing cells. Representative examples of respective phenotypes are shown on the right. Scale bar is 10 mm. N = 276, 266, and 281 cells analyzed for MiD51 WT, Y185A, R342A, respectively. **** denotes p value of ≤ 0.0001 by ANOVA (Dunnett’s multiple comparisons) test. n.s. = not significant (p value > 0.05). **c,** Western blot showing expression levels of GFP-fusion constructs, versus endogenous MiD51 using anti-MiD51 (green). Actin and tubulin levels in red. **d,** Box and whiskers plot of average MiD51 puncta size for MiD51-WT, MiD51-Y185A and MiD51-R342A. Number of cells analyzed per condition denoted by n. **** indicates a p value of ≤ 0.0001 and n.s. (not significant) corresponds to p > 0.05 by ANOVA (Dunnett’s multiple comparisons) test. **e,** Box and whiskers plot of % mitochondrial area covered by MiD51 staining, for MiD51-WT, MiD51-Y185A and MiD51-R342A. Number of cells analyzed per condition denoted by n. **** indicates a p-value of ≤ 0.0001 by ANOVA (Dunnett’s multiple comparisons) test. Bars in panels b, d and e represent standard error of the mean.

We have previously shown that actin filaments synergize with MFF in stimulating DRP1, with actin filaments decreasing the concentration of MFF needed for optimal stimulation^24,26^. We tested the possibility that MiD49 oligomers might act in a similar manner to actin. Indeed, an MiD49 concentration that causes sub-optimal DRP1 activation alone (250 nM) allows further DRP1 activation by low concentrations of MFF (EC_50_ 1 µM, **Fig. 2f, Extended Data Fig. 5b**), as opposed to the situation for MFF alone, in which 100 µM MFF is required for full DRP1 activation (**Extended Data Fig. 5c**). In contrast, MiD49 monomer reduces the effectiveness of MFF in DRP1 activation (**Fig. 2f, Extended Data Fig. 5b, c**). We tested whether MiD49 oligomers, like MFF, had the ability to synergize with actin filaments. However, varying concentrations of actin filaments do not change the effect of MiD49 monomers or oligomers on DRP1 activity (**Fig. 2g**). In addition, MiD49 monomers inhibit the effect of actin filaments on DRP1 GTPase activity (**Fig. 2g**), with an IC_50_ below 500 nM (**Fig. 2h**).

These results suggest that LCACA-induced oligomerization converts MiD49 into a DRP1-activating protein. MiD49 oligomers are synergistic with MFF for DRP1 activation, in a similar manner to the synergism between actin filaments and MFF. In contrast, MiD49 and actin filaments do not synergize. In fact, MiD49 monomers eliminate the activating effect of actin filaments on DRP1 activity, suggesting that the two are competitive for DRP1 binding.

### MiD51 displays acyl-CoA induced oligomerization

We next tested the ability of LCACA to induce oligomerization of the cytoplasmic region of MiD51 (**Extended Data Fig. 1a,b**). Similar to MiD49, MiD51 migration on BNGE is slower in the presence of acyl-CoA of 12 carbons or longer, but not with octanoyl-CoA, acetyl-CoA, malonyl-CoA, CoA alone, palmitic acid, lyso-PA (**Fig. 3a**), or a series of purine nucleotides (**Extended Data Fig. 6a**). By size-exclusion chromatography, palmitoyl-CoA causes a shift in MiD51 to an apparent oligomer, with the size depending on the palmitoyl-CoA:MiD51 ratio in the oligomerization reaction. Lower palmitoyl-CoA concentrations cause migration in the void volume (**Extended Data Fig. 6b**) and higher concentrations result in an oligomer that is resolved in the column (**Fig. 3b**). By velocity analytical ultracentrifugation, this resolved oligomer sediments at 51 S (**Fig. 3c**), and has an apparent mass of 2450 kDa (**Extended Data Fig. 6c**), suggesting oligomers averaging 68 subunits. In contrast, the MiD51 peak in the absence of palmitoyl-CoA elutes similarly to the ovalbumin marker by size-exclusion chromatography (**Fig. 3b**) and sediments at 2.9 S (**Fig. 3c**) with a calculated mass of 38.3 kDa (**Extended Data Fig. 6c**), suggestive of a monomer. By negative stain electron microscopy, the oligomer displays a range of particle sizes, generally smaller than those for MiD49 oligomers (**Fig. 3d, Extended Data Fig. 2**). We assessed palmitoyl-CoA:MiD51 ratio across the oligomer peak, and found an approximate 1:1 correspondence (**Fig. 3e**). These results suggest that the increase in MiD51 size is due to LCACA-induced oligomerization of MiD51.

**Fig. 6.**
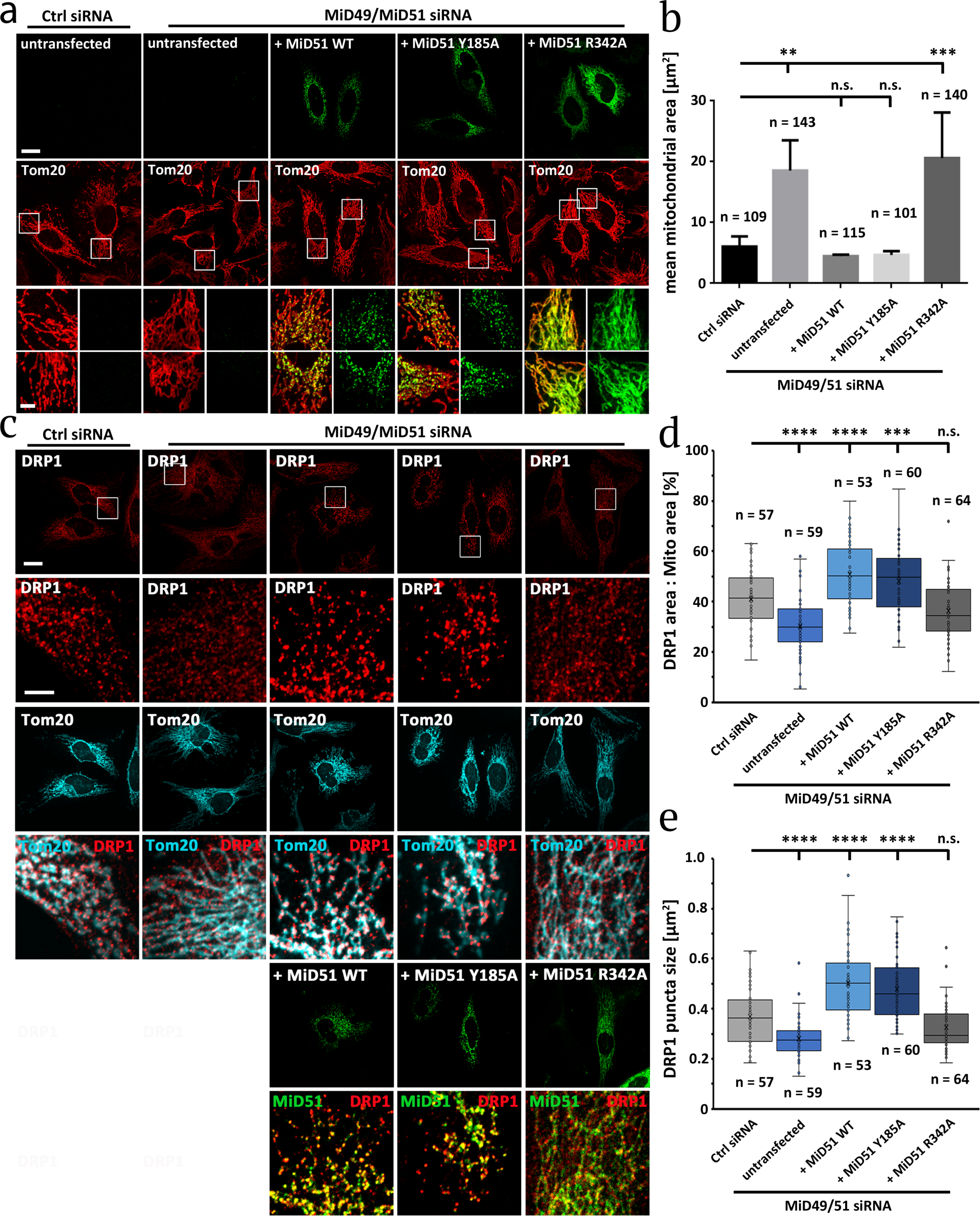
Effect of LCACA binding defect on ability of MiD51 re-expression to rescue mitochondrial elongation caused by MiD49/51 suppression. **a,** C-terminal GFP fusion constructs of MiD51-WT, MiD51-Y185A or MiD51-R342A were transiently transfected into HeLa cells that had been previously treated with siRNAs against MiD49 and MiD51. Cells were then fixed and stained for mitochondria (Tom20). Cells displaying low GFP levels were imaged. Boxed regions denote insets shown in zoom to the right of full images. Zoom images either display MiD51-GFP signal alone or as corresponding merge image with Tom20. Scale bars, 20 mm in overview images and 5 mm in inset images. **b,** Graph of mean mitochondrial area, determined as described in Methods, under the indicated conditions. N represents the number of cells analyzed per respective condition. ** and *** indicate p values ≤ 0.01 and ≤ 0.001 by ANOVA (Dunnett’s multiple comparisons) test. n.s. = not significant (p value > 0.05). Statistical test was performed on the average values of four independent experiments. **c,** Drp1 distribution on mitochondria in control versus MiD49/51 siRNA-treated HeLa cells either without or with re-expression of the indicated MiD51-GFP construct. Cells fixed and stained for Drp1 and Tom20. Scale bars, 20 mm in overview images and 5 mm in zoom images. **d,** Quantification of % mitochondrial area covered by Drp1 for the indicated conditions. **** and *** indicate p values ≤ 0.0001 and ≤ 0.001 by ANOVA (Dunnett’s multiple comparisons) test. n.s. = not significant (p value > 0.05). N represents the number of cells analyzed per condition. **e,** Quantification of Drp1 puncta size for the indicated conditions. **** indicate a p value of ≤ 0.0001 by ANOVA (Dunnett’s multiple comparisons) test. n.s. = not significant (p value > 0.05). N represents the number of cells analyzed per condition. Bars in b, d and e represent standard error of the mean.

MiD51 has been previously shown to bind ADP^19,20^. We find that ADP does not induce MiD51 oligomerization (**Extended Data Fig. 6d**). ADP does decrease palmitoyl-CoA induced MiD51 oligomerization at higher concentrations, whereas CoA or acetyl-CoA have less effect (**Extended Data Fig. 6d**). This result suggests that ADP and palmitoyl-CoA compete for MiD51 binding. Even at 0.5 mM ADP, however, significant MiD51 still oligomerizes.

We took advantage of a previously used assay to examine MiD51 binding specificity in more detail. In this assay^20^, MiD51 is mixed with fluorescent MANT-ADP, and the change in MANT-ADP fluorescence is monitored, with binding correlating with higher fluorescence intensity. Similar to the past study, MiD51 binds MANT-ADP with an apparent K_d_ of 0.65 µM, while MiD49 displays no detectable MANT-ADP binding (**Extended Data Fig. 6e**). We then assessed MiD51 binding to potential ligands using a competition assay in which increasing competitor ligand is mixed with fixed concentrations of MANT-ADP and MiD51. Palmitoyl-CoA causes a concentration-dependent decrease in MANT-ADP fluorescence back to the level of MANT-ADP alone, with an EC_50_ of 2 µM (**Fig. 3f**), suggesting competition for the same binding site on MiD51. Using this assay, we screened several molecules at fixed concentration (20 µM) for competition with MANT-ADP. LCACAs (stearoyl, oleoyl, palmitoyl, myristoyl) compete efficiently, whereas lauroyl-CoA, octanoyl-CoA, malonoyl-CoA, acetyl-CoA, CoA alone, palmitoyl-carnitine, and lyso-phosphatidic acid display no competition (**Fig. 3g**). Interestingly, ADP is a poor competitor for MANT-ADP (**Fig. 3f**), suggesting that the hydrophobic MANT group contributes significantly to the affinity of MiD51 for MANT-ADP in a similar manner to the fatty acid tail of acyl-CoA.

As with MiD49, oligomerized MiD51 activates DRP1 GTPase activity, whereas monomeric MiD51 does not (**Extended Data Fig. 6f**). Also similar to MiD49, MiD51 oligomers activate DRP1 in a bi-phasic manner, with an optimal concentration at 1 µM (**Fig. 3h**). The optimal concentration increases if DRP1 concentration is increased (**Fig. 3i**), supporting the model of increased DRP1 density on MiD51 oligomers to allow GTPase domain interaction (**Fig. 2d**). In addition, MiD51 oligomers synergize with MFF for DRP1 activation, with the EC_50_ of MFF being 2 µM in the presence of MiD51 oligomers, which is over 10-fold lower than for MFF alone (**Fig. 3j, Extended Data Fig. 6g**). In contrast, the same concentration of MiD51 monomers reduces MFF-mediated DRP1 activation (**Fig. 3j, Extended Data Fig. 6g**). Similar to MiD49, MiD51 displays no ability to synergize with actin filaments in DRP1 activation (**Extended Data Fig. 6h**), and MiD51 monomers are potent inhibitors of actin-stimulated DRP1 GTPase activity (**Fig. 3k**).

These results suggest that, similar to MiD49, MiD51 binding to LCACA induces oligomerization, and these oligomers are capable of DRP1 activation. While MiD51 is capable of binding ADP, LCACAs are preferred ligands, presumably constituting the physiological ligands in cells. MiD51 oligomers synergize with MFF in DRP1 activation, while MiD51 monomers inhibit both MFF- and actin-mediated DRP1 activation.

### Mutation of R342 in MiD51 reduces LCACA-induced oligomerization

We sought to design an MiD51 mutant deficient in LCACA binding, using published structural information as a guide ^19,20^. We made mutations to two residues protruding from either side of MiD51’s pocket, Y185A and R342A. The R342 side chain is in close proximity to the alpha phosphate of both ADP and GDP, whereas Y185 is in the vicinity of the purine ring (**Fig. 4a**). The cytoplasmic regions of both MiD51-R342A and MiD51-Y185A express and purify in bacteria similar to WT MiD51 (**Extended Data Fig. 1b**), eluting from SEC as apparent monomers (**Fig. 4b**). MiD51-R342A displays an apparent inability to bind MANT-ADP, whereas MiD51-Y185A displays binding similar to WT (K_d_ WT 0.27 µM, K_d_ Y185A 0.9 µM, **Fig. 4c**). By BNGE, MiD51-R342A displays a reduced ability to oligomerize in response to palmitoyl-CoA (**Fig. 4d**), quantified by loss of the MiD51 monomer band with increasing palmitoyl-CoA (**Fig. 4e**). Similarly, a lower amount of oligomer peak is recovered from SEC for the R342A mutant upon incubation with palmitoyl-CoA (**Fig. 4f**). Interestingly, the MiD51-R342A oligomer maintains the ability to activate DRP1 GTPase activity (**Fig. 4g**) suggesting that, although MiD51-R342A has reduced affinity for LCACA, it maintains DRP1 binding and activation once oligomerized.

### Acyl-CoA binding mutants display reduced oligomerization and mitochondrial phenotypes in cells

At low expression levels, MiD proteins display a punctate appearance^15,17^, suggestive of oligomerization. At higher expression, MiD distribution is uniform and causes mitochondrial elongation by its ability to sequester DRP1 ^11,15,17^. We found similar effects with GFP-fusions of MiD49 and MiD51 (**Fig. 5a, Extended Data Fig. 7a**). Live-cell imaging suggests that, for both MiD49 and MiD51, the puncta appear stable on the mitochondria, despite considerable fluctuation of the mitochondria themselves (**Movies 1, 2**).

**Fig. 7.**
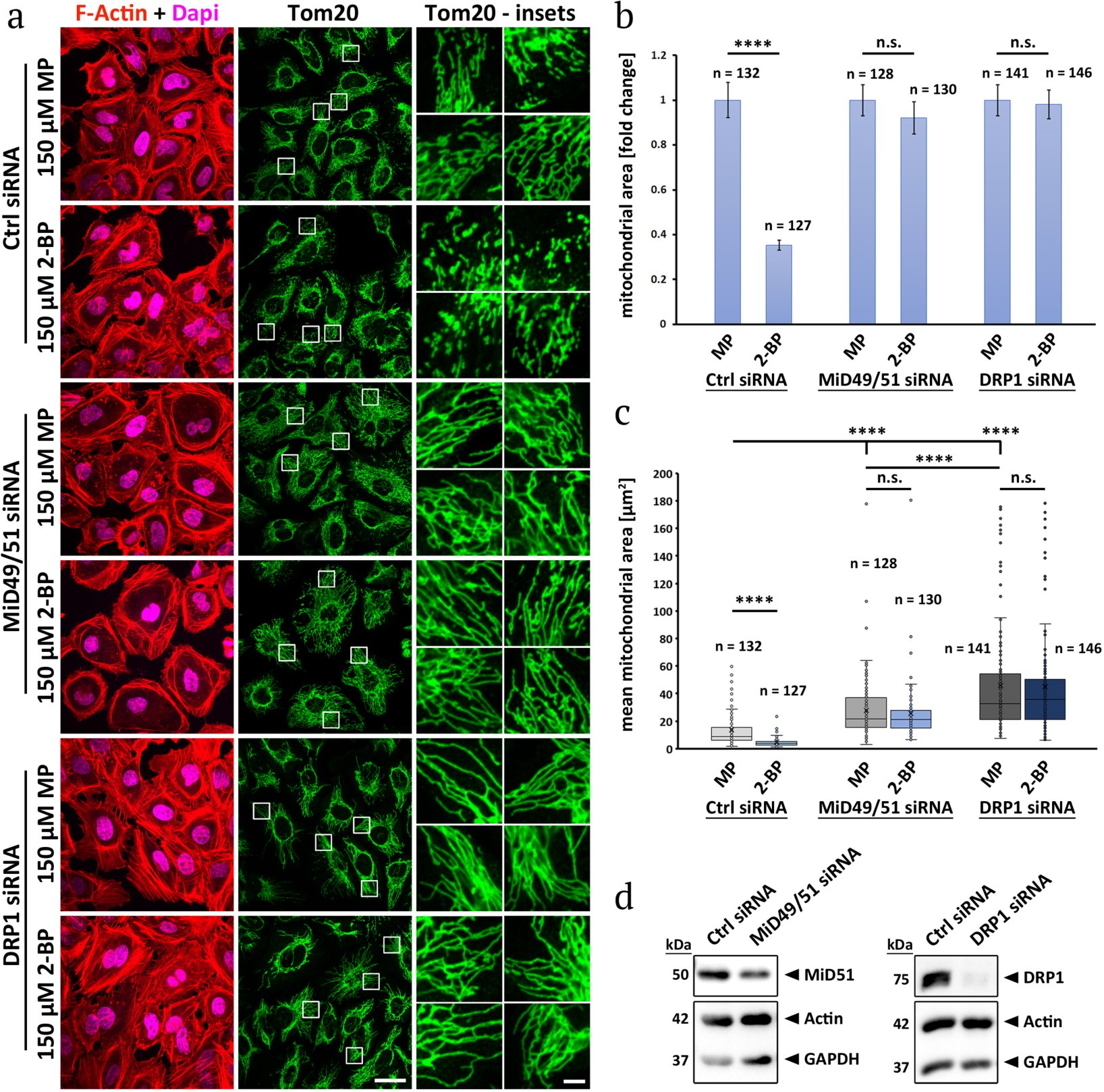
2-bromopalmitate induced mitochondrial fission is MiD-dependent. **a,** Fixed-cell fluorescence micrographs of HeLa cells transfected with the indicated siRNAs, treated for 1-hr with either 2-BP or MP and stained for actin (TRITC-phalloidin, red) with Dapi (magenta) and mitochondria (Tom20, green). **b,** Quantification of relative mitochondrial lengths between the MP and 2-BP treatments. Data normalized to the MP treatment. Corresponding non-normalized data are in Panel C. **** indicates p value ≤ 0.0001 by unpaired Student’s t-test. n.s. = not significant (p value > 0.05). N represents the number of cells analyzed per condition. **c,** Quantification of mean mitochondrial area between the MP and 2-BP treatments. Corresponding normalized data are in Panel B. **** indicates p value ≤ 0.0001 by unpaired Student’s t-test. n.s. = not significant (p value > 0.05). N represents the number of cells analyzed per condition. Mean values are as follows: Ctrl siRNA/MP = 13.46; Ctrl siRNA/2-BP = 4.75; MiD49/51 siRNA/MP = 27.64; MiD49/51 siRNA/2-BP = 25.44; DRP1 siRNA/MP = 45.88; DRP1 siRNA/2-BP = 44.97. **d,** Western blot of HeLa cells knocked down for either MiD49/MiD51 or DRP1, confirming the efficiency of the used siRNAs. Actin and GAPDH serve as loading controls. Bars in b and c represent standard error of the mean.

Identification of an MiD51 mutant defective in LCACA binding provided an opportunity to test the importance of LCACA binding to MiD function in cells. We expressed GFP-fusions of WT, Y185A or R342A MiD51 in HeLa cells and evaluated their ability to form puncta at low expression level, as well as their effect on mitochondria at high expression level. Low level expression of either MiD51-WT or MiD51-Y185A results in punctate GFP accumulation on mitochondria, in contrast to the even distribution of MiD51-R342A (**Fig. 5a,b, Movies 2-4**). The expression levels of these constructs are similar, and lower than endogenous MiD51 levels (**Fig. 5c**). We quantified MiD51 puncta by examining the average size of each MiD51 particle and the % of mitochondrial area covered by MiD51 staining (examples in **Extended Data Fig. 7b**), with lower numbers denoting puncta. Both average particle size and % mitochondrial area are significantly lower for MiD51-WT and MiD51-Y185A than for MiD51-R342A (**Fig. 5d,e**). At higher expression, MiD51-WT and MiD51-Y185A are evenly distributed along mitochondria and cause mitochondrial elongation, while MiD51-R342A causes a collapsed mitochondrial phenotype, which could be due to dominant negative effects of this mutant on the endogenous MiD51 and/or MiD49 proteins (**Extended Data Fig. 7c,d**).

We next tested whether MiD51-WT, MiD51-Y185A or MiD51-R342A could rescue phenotypes caused by siRNA-mediated double knock-down (KD) of MiD49 and MiD51. MiD49/51 KD in HeLa cells results in significant mitochondrial elongation (**Fig. 6a**). Expression of MiD51-WT or MiD-Y185A results in substantial rescue of the mitochondrial elongation phenotype, while expression of MiD51-R342A does not (**Fig. 6a,b**). In addition, MiD51-R342A appears diffuse on mitochondria in the MiD49/51 KD cells (**Fig. 6a**), similar to its distribution in WT HeLa cells (**Fig. 5a).**

Knock-down of MiD49/51 also results in a decrease in the punctate appearance of DRP1 on mitochondria (**Fig. 6c**), similar to results from other studies^13–15^. We quantified these effects either by % mitochondrial area covered by DRP1 (**Fig. 6d**) or the size of DRP1 puncta (**Fig. 6e**). Re-expression of MiD51-WT or MiD51-Y185A results in a recovery of DRP1 puncta to larger sizes and mitochondrial coverage than in control cells (**Fig. 6c-e**), suggesting enhanced DRP1 oligomerization. In contrast, expression of MiD51-R342A does not cause recovery of DRP1 puncta size or overall mitochondrial area covered by DRP1 (**Fig. 6c-e**). The combination of these results suggest that LCACA binding is required for MiD51-mediated DRP1 recruitment and mitochondrial fission.

### Increased cellular LCACA causes increased MiD-dependent mitochondrial fission

To test the involvement of LCACA in MiD-mediated mitochondrial fission, we treated cells with 2-bromopalmitate (2-BP), a palmitate analogue that gets converted to 2-bromopalmitoyl-CoA that is slow in subsequent processing ^31,32^. We treated HeLa cells for 1-hr with either 150 µM 2-BP or methyl-palmitate (MP) as a negative control ^32^, and evaluated the mitochondrial phenotype. In HeLa WT cells, a 1-hr treatment with 2-BP causes a 2.8-fold decrease in mitochondrial area after 1 h (**Fig. 7a-c**), as well as an increase in mitochondrial DRP1 accumulation (**Extended Data Fig. 8**) when compared to control (MP-treated) cells. DRP1 KD causes mitochondrial area to increase 3.4-fold in control cells, and eliminates the 2-BP mediated decrease, suggesting that the change in mitochondrial length is due to increased fission (**Fig. 7a-d**). Similarly, MiD49/51 KD causes mitochondrial area to increase 2-fold in control cells, and eliminates the 2-BP-induced mitochondrial shortening (**Fig. 7a-d**). These results suggest that 2-BP mediated mitochondrial fission occurs through increased acyl-CoA levels, activating MiD49/51.

**Fig. 8.**
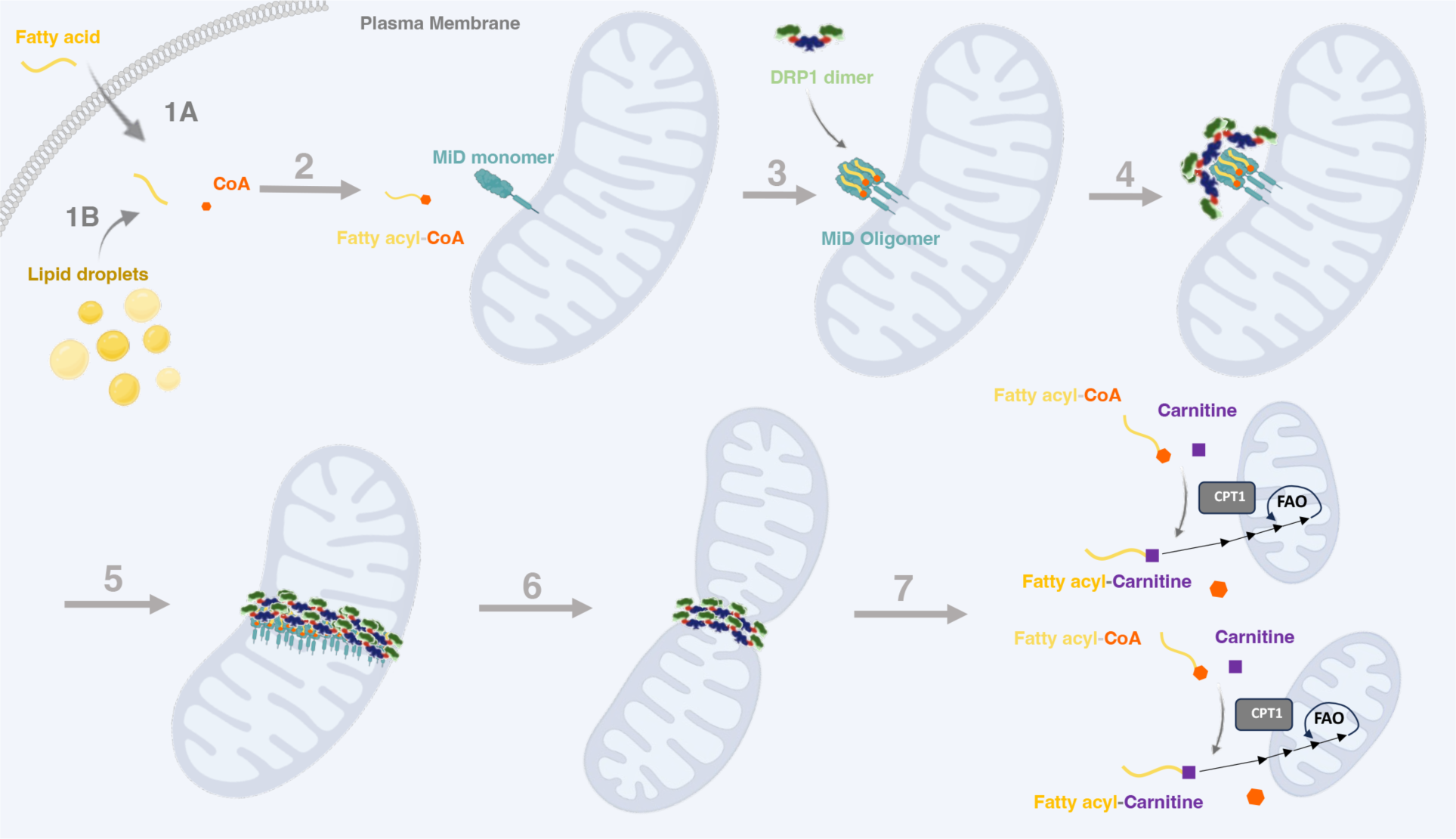
Model for MiD function in activation of fatty acid oxidation. This model is developed from results in this paper and from (Ngo *et* al, 2023). <u>Step 1</u>. Long-chain fatty acid enters cytoplasm from the extracellular milieu (1A) or from intracellular lipid droplets (1B). <u>Step 2</u>. Fatty acid is coupled to CoA through fatty acyl-CoA synthetase. <u>Step 3</u>. Fatty acyl-CoA binds to MiD49 and/or MiD51 monomers on the outer mitochondrial membrane (OMM), inducing their oligomerization. <u>Step 4</u>. Oligomerized MiD initiates assembly of active DRP1 oligomers on the OMM. <u>Step 5</u>. DRP1 oligomerization continues on the OMM to create a ring around the mitochondrion. <u>Step 6</u>. DRP1-mediated constriction of the OMM leads to mitochondrial fission. <u>Step 7</u>. Mitochondrial fission causes increased carnitine-palmitoyl transferase 1 (CPT1) activity, increasing fatty acid import into the mitochondrial matrix and subsequent fatty acid oxidation (FAO).

## Discussion

We report that long chain fatty acyl CoAs (LCACAs) are ligands for the mitochondrial DRP1 receptors MiD49 and MiD51, triggering oligomerization which in turn enables activation of the DRP1 GTPase. LCACA-oligomerized MiD can synergize with MFF in activating DRP1, but is not synergistic with actin. A mutant compromising LCACA binding reduces cellular MiD51 oligomerization and its ability to enhance mitochondrial fission. Cellular treatment with 2-bromopalmitic acid, which increases acyl-CoA levels, stimulates mitochondrial fission in an MiD-dependent manner.

The identification of LCACAs as MiD ligands answers an outstanding question concerning these proteins. Both MiD49 and MiD51 contain large interior pockets similar to that of cGAS, but no ligand had been identified for MiD49, whereas the ligands identified for MiD51 (ADP or GDP) do not result in a dramatic change in its ability to activate DRP1^19,20^. We find that the affinity of MiD51 for palmitoyl-CoA is significantly higher than for ADP, suggesting that LCACAs likely out-compete ADP for MiD51 binding in cells. The demonstration that LCACAs bind with 1:1 stoichiometry to MiDs and cause a substantial increase in DRP1 activation suggests that these are physiological ligands. In addition, the fact that palmitoyl-CoA concentrations below its critical micelle concentration (measured here to be 47 µM) can induce MiD oligomerization suggests that MiDs are not simply aggregating around micelles. Furthermore, LCACAs are preferred over other possible ligands such as lyso-phosphatidic acid, acetyl-CoA, malonyl-CoA, or a variety of nucleotides.

The activation mechanism induced by LCACA is intriguing. In this study and previous studies^19,20^, the cytoplasmic regions of MiD49 or MiD51 cannot activate DRP1, and even display an inhibitory effect on GTPase activity. We show that LCACA-induced MiD oligomers activate DRP1. This activation is presumably induced by bringing GTPase domains in close proximity, which is a common activation mechanism for dynamin proteins in general^6–8^. The fact that MiDs display bi-phasic concentration effects on DRP1 activation, with activation reaching a peak and then declining at higher MiD oligomer concentrations, further suggests activation through inducing GTPase domain proximity.

DRP1 recruitment to and activation by oligomeric ‘receptors’ occurs in two other contexts: with MFF, which requires oligomerization through its coiled-coil^24^; and with actin filaments^24,26^. A recent paper revealed the structure of ‘cofilaments’ assembled by DRP1 and MiD49 in the presence of GTP or non-hydrolyzable GTP analogues^33^. In this structure, monomeric MiD49 is bound around a core of DRP1, and GTP hydrolysis causes MiD49 release. It is unclear how the assembly of MiD49 oligomers might alter this interaction.

Another intriguing aspect of MiD effects on DRP1 is the ability of oligomerized MiD49 or MiD51 to synergize with MFF in DRP1 activation. MFF alone is a poor DRP1 activator, requiring high concentrations to stimulate DRP1 activity. Addition of a low concentration of MiD49 or MiD51 oligomers reduces the concentration of MFF required for DRP1 activation. We have observed a similar effect for actin filaments^24^.

These results raise a model whereby either activation of MiD proteins (by increased LCACA) or polymerization of specific actin-based structures (through the polymerization factors INF2 or Spire1C^27,34^) might serve as initiating signals for DRP1 recruitment, with MFF working downstream. A previous study suggests that MFF preferentially binds DRP1 oligomers^35^, supporting the model. This model might also suggest that MiDs and actin operate in distinct mitochondrial fission events, which is further supported by the lack of synergy between oligomerized MiDs and actin, as well as the potent inhibition of actin-mediated DRP1 activation by monomeric MiDs. While the possibility of actin and MiDs operating in distinct pathways has not been tested directly, a recent publication reveals two mechanistically distinct pathways to DRP1-dependent mitochondrial fission^36^, one which is actin-dependent and one which is actin-independent. Interestingly, deletion of MiD49 and MiD51 partially reduces both types of fission in this study, suggesting that additional factors might be at play.

Another question pertains to the pools of LCACA that stimulate MiD activity in cells. LCACAs are involved in both anabolic (phospholipid and triacylglycerol synthesis) and catabolic (beta-oxidation) processes, in addition to being required for protein myristoylation and palmitoylation. In principle, processes involving LCACA on the mitochondrial surface, such as fatty acid import into mitochondria for beta-oxidation, would be the most likely candidates for MiD regulation. For fatty acid import, long chain fatty acids are first coupled to CoA in the cytoplasm, then converted to acyl-carnitine by carnitine O-palmitoyltransferase 1 (CPT1) on the outer mitochondrial membrane^37^.

A recently published paper has established a correlation between increased mitochondrial fission and increased fatty acid oxidation, with the fission-activated step being CPT1^3^. A possible mechanism for fission-mediated CPT1 activation is through increased membrane curvature, facilitating intra-molecular interactions^38,39^. The results presented here, in combination with this recent publication, lead to a model whereby increased cytoplasmic fatty acid leads to increased fatty acyl-CoA, activating MiD proteins to trigger mitochondrial fission. Fission leads to CPT1 activation, increased fatty acid import and beta-oxidation (**Fig. 8**), which could be used for ATP production (hepatocytes, pancreatic beta-cells, and oxphos-dependent B lymphoma cells ^3^) or for heat generation (brown adipocytes ^29^). Interestingly, a recent publication suggests that CPT1 might have a reciprocal effect, enhancing mitochondrial fission through succinylation-mediated stabilization of MFF^40^.

MiD proteins are only found in metazoans, so it is unclear whether LCACA influences mitochondrial fission in non-metazoans. Interestingly, knocking out cytoplasmic acyl-CoA binding protein in *Schizosaccharomyces pombe* causes DRP1-dependent mitochondrial fission ^41^, implying that increasing levels of free acyl-CoA induces mitochondrial fission. The proteins mediating this effect remain to be elucidated.

The regulation of MiD49/51 adds to a growing list of LCACA regulatory functions. LCACAs are allosteric inhibitors of acetyl-CoA carboxylase (ACC), with this inhibition simultaneously decreasing fatty acid synthesis and relieving inhibition of CPT1 ^32^. Interestingly, the basis for this inhibition might be in regulating ACC’s oligomeric state^42^. Recently, LCACAs have also been shown to act as allosteric activators AMP-dependent kinase beta1, further inhibiting ACC ^32^. All of these effects lead to more efficient mitochondrial fatty oxidation, and could act in a concerted fashion toward these goals.

## METHODS

### Plasmids/siRNA

For bacterial expression, MiD49Δ1-124 (mouse amino acids 125-454, UniProt ID Q5NCS9) was inserted into a modified pGEX-KT vector by BamH1 and EcoR1 sites, in which a GST-thrombin site-6xHis-TEV tag is in front of multiple cloning site (MCS). MiD51Δ1-133 (human amino acids 134-463, UniProt ID Q9NQG6) was inserted into another modified pGEX-KT vector by BamH1 and EcoR1 sites, where a GST-thrombin site-6xHis-HRV3C-6xHis sequence is in front of MCS. Quick Change mutagenesis was performed to make MiD49 or MiD51 mutants. Full-length of human DRP1 isoform 3 (NP_055681.2, UniProt ID O00429-4) and truncated human MFF isoform 4 (UniProt ID Q9GZY8-4) (MFF-ΔTM) have been described previously ^14,26^. For cellular assays, full length MiD49/51 and corresponding mutants were inserted into a modified GFP-N1 vector, in which the MCS is followed by HRV3C-GFP-2xStrep-tag. Mito-plum plasmid was obtained from Addgene (#55988). The following oligonucleotides used for siRNA-mediated protein silencing were synthesized by Integrated DNA Technologies: MiD49 (3’UTR Exon 4): 5’-AUUCUGACUUUGAAGCCUGUUAAGA-3’; MiD51 (3’UTR Exon 6): 5′-GAAGAGCUGUGAUAGCAUGUUUCAA-3’; DRP1 (CDS Exon 8): 5’-GCCAGCTAGATATTAACAACAAG AA-3′; silencer negative control (IDT) was 5′-CGUUAAUCGCGUAUAAUACGCGUAU-3′.

### Protein Expression, Purification

MiD49 and MiD51 was expressed in One Shot BL21 Star (DE3) *Escherichia coli* (C6010-03; Life Technologies, Carlsbad, CA) in LB broth, induced by isopropyl-β-D-thiogalactoside (IPTG) at 16 °C for 16 h when OD600 reached 1.5. Cell pellets were resuspended in MiD lysis buffer (25 mM 4-(2-hydroxyethyl)-1-piperazineethanesulfonic acid [Hepes], pH 7.4, 500 mM NaCl, 1 mM dithiothreitol [DTT], 2 µg/ml leupeptin, 10 µg/ml aprotinin, 2 µg/ml pepstatin A, 2 mM benzamidine, 1 µg/ml calpain inhibitor I [ALLN], and 1 µg/ml calpeptin) and lysed using a high-pressure homogenizer (M-110L Microfluidizer Processor; Microfluidics, Newton, MA). The lysate was cleared by centrifugation at 40,000 rpm (type 45 Ti rotor; Beckman, Brea, CA) for 1 hour at 4°C and then was loaded onto Pierce™ Glutathione Agarose (16101; Thermofisher) by gravity flow. The column was washed with 20 column volumes (CV) of lysis buffer without protease inhibitors. To elute MiD49/51, 1 unit/µL thrombin (T4648, Sigma-Aldrich) or 0.01 mg/ml HRV3C protease in lysis buffer without protease inhibitors was added for 16 hours at 4°C. The protein eluate was captured by HiTrap IMAC column (17-5248-01, GE Healthcare, Chicago, IL) and eluted by IMAC-B buffer (50 mM Tris-HCl pH 7.5, 0.1 M NaCl, 500 mM imidazole). The His-trap protein eluate was further purified by size exclusion chromatography on Superdex200 (GE Biosciences, Piscataway, NJ) with S200 buffer (20 mM Hepes, pH 7.4, 65 mM KCl, 2 mM MgCl_2_, 1 mM DTT, 0.5 mM ethylene glycol tetraacetic acid [EGTA]), spin concentrated (UFC903024, EMD Millipore Corporation, Burlington, MA), frozen in liquid nitrogen, and stored at −80 °C.

DRP1 was expressed and purified as previously described with modifications^26^. Briefly, DRP1 construct was expressed in One Shot BL21 Star (DE3) *Escherichia coli* in LB broth, induced by isopropyl-β-D-thiogalactoside (IPTG) at 16 °C for 16 hours when OD600 reached to 1.5. Cell pellets were resuspended in lysis buffer (100 mM Tris-Cl, pH 8.0, 500 mM NaCl, 1 mM dithiothreitol [DTT], 1 mM Ethylenediaminetetraacetic acid [EDTA], 2 µg/ml leupeptin, 10 µg/ml aprotinin, 2 µg/ml pepstatin A, 2 mM benzamidine, 1 µg/ml ALLN, and 1 µg/ml calpeptin) and lysed using a high-pressure homogenizer. The lysate was cleared by centrifugation at 40,000 rpm in Ti-45 rotor for 1 hour at 4°C. Avidin (20 µg/ml; PI-21128; Thermo Fisher Scientific, Waltham, MA) was added to the supernatant, and then was loaded onto Strep-Tactin Superflow resin (2-1206-025; IBA, Göttingen, Germany) by gravity flow. The column was washed with 20 column volumes (CV) of lysis buffer without protease inhibitors. To elute DRP1, 0.01 mg/ml HRV3C protease in lysis buffer without protease inhibitors was added for 16 hours at 4°C. The Strep-Tactin Superflow eluate was further purified by size exclusion chromatography on Superdex200 with DRP1-S200 buffer (20 mM HEPES pH 7.5, 150 mM KCl, 2 mM MgCl_2_, 1 mM DTT, 0.5 mM EGTA), spin concentrated, frozen in liquid nitrogen, and stored at −80 °C.

MFF-ΔTM was expressed in Rosetta^TM^2 BL21-(DE3) *Escherichia coli* (71400; EMD Millipore Corporation, Burlington, MA) in LB broth, induced by 1M IPTG at 30 °C for 4 h when OD600 reached to 1.5. Cell pellets were resuspended in lysis buffer (50 mM Tris-HCl, pH 7.5, 500 mM NaCl, 20 mM imidazole, pH 7.5, 1 mM DTT, 1 mM EDTA, 2 µg/ml leupeptin, 10 µg/ml aprotinin, 2 µg/ml pepstatin A, 2 mM benzamidine, 1 µg/ml ALLN, and 1 µg/ml calpeptin) and lysed using M-110 microfluidizer processor. The lysate was cleared by centrifugation at 40, 000 rpm in Ti45 for 40 minutes at 4°C, the supernatant was saved. Affinity capture was performed using FPLC and a HiTrap IMAC column (17-5248-01, GE Healthcare, Chicago, IL) equilibrated with IMAC-A buffer (50 mM Tris-HCl pH 7.5, 0.1 M NaCl, 20 mM imidazole). Cleared lysate was loaded onto the column with a rate of 3 mL/min and washed to baseline with IMAC-A. MFF was eluted from the column with gradient step washes by IMAC-B buffer (50 mM Tris-HCl pH 7.5, 0.1 M NaCl, 500 mM imidazole): step1 10% IMAC-B for 5CV, step2 20% IMAC-B for 5CV, step3 100% for 5CV. Fractions from step3 were pooled and diluted 10-fold in ion exchange (IEX)-A buffer (50 mM Tris-HCl pH 7.5, 1 mM DTT). Diluted fractions were loaded onto a HiTrap Q anion exchange column (54816, EMD Millipore Corporation, Burlington, MA). The column was washed to baseline with IEX-A and MFF was eluted by IEX-B buffer (50 mM Tris-HCl pH 7.5, 1 M NaCl, 1 mM DTT) with a step gradient: step1 10% 5CV, linear 10-50% 30CV followed by linear 50-100% 5CV. Peak MFF fractions were concentrated by reloading onto the HiTrap IMAC column and eluted with 100% IMAC-B step wash. MFF fractions were pooled and further purified by size exclusion chromatography on Superdex200 with S200 buffer (20 mM HEPES, pH 7.4; 2 mM MgCl_2_, 0.5 mM EGTA, 65 mM KCl, 1 mM DTT), spin concentrated (UFC903024, EMD Millipore Corporation, Burlington, MA), aliquots were frozen in liquid nitrogen, and stored at −80 °C.

Rabbit skeletal muscle actin was extracted from acetone powder as previously described ^43^, and further gel-filtered on Superdex 75 16/60 columns (GE Healthcare). Actin was stored in G buffer (2 mM Tris, pH 8.0, 0.5 mM DTT, 0.2 mM ATP, 0.1 mM CaCl_2_, and 0.01% NaN_3_) at 4°C.

### Actin preparation for biochemical assays

For high-speed pelleting assay, actin filaments were polymerized from 20 µM monomers for 3 h at 23 °C by addition of a 10x stock of polymerization buffer (200 mM HEPES, pH 7.4, 650 mM KCl, 10 mM MgCl_2_, 10 mM EGTA) to a final 1x concentration. For GTPase assay, actin monomers in G-buffer were incubated with AG1-X2 100–200 mesh anion exchange resin (Dowex; 1401241; Bio-Rad) at 4 °C for 5 min to remove ATP, followed by low-speed centrifugation. 20 µM actin filaments were polymerized as described before. To maintain ionic strength across all samples, an actin blank was prepared in parallel using G-buffer in place of actin monomers and used to dilute actin filaments as needed for each sample. DRP1 was diluted in MEHD buffer (20 mM HEPES, pH 7.4, 2 mM MgCl_2_, 0.5 mM EGTA, 1 mM DTT) to adjust the ionic strength to the same as S200 buffer before biochemical assays.

### Size exclusion Chromatography assays

MiD49Δ1-124 and MiD51Δ1-133 oligomeric distribution was determined by Superose 6 increase 10/300 GL SEC column (GE Biosciences) in S200 buffer (20 mM HEPES, pH7.4, 65 mM KCl, 2 mM MgCl_2_, 0.5 mM EGTA, 1 mM DTT). Protein at varying concentration was loaded onto the column in a total volume of 500 µL and gel-filtered with a flow rate of 0.4 mL/min.

### Purine-containing ligand screening and Blue-Native PAGE

Purine-containing ligands were incubated with MiD49 or MiD51 at 37 °C for 1 hour before BN-PAGE analysis. Ligands include: NADP+ (Sigma-Aldrich, 077K7000), NAD+ (Sigma-Aldrich, N1636), NADH (Sigma-Aldrich, 10107735001), c-di-AMP (Sigma-Aldrich, A3262), 2’3’-GAMP (Sigma-Aldrich, 1229), AMP (Sigma-Aldrich, 01930), ADP(Sigma-Aldrich, 01897), ATP (Sigma-Aldrich, A2383), GMP (Sigma-Aldrich, G8377), GDP (Sigma-Aldrich, G7127), GTP (Sigma-Aldrich, G8877), cAMP (Sigma-Aldrich, 1231), dNTP (New England Biolabs, N0447S), hypoxanthine (Sigma-Aldrich, H9377), stearoyl-CoA (Sigma-Aldrich, S0802), oleoyl-CoA (Sigma-Aldrich, O1012), palmitoyl-CoA (Sigma-Aldrich, P9716), myristoyl-CoA (Sigma-Aldrich, M4414), lauoryl-CoA (Sigma-Aldrich, L2659), octanoyl-CoA (Sigma-Aldrich, O6877), malonoyl-CoA (Sigma-Aldrich, 4263), acetyl-CoA (Sigma-Aldrich, A2056), coenzyme A (Sigma-Aldrich, C4282), lyso-PA (Avanti Polar Lipids, 857127P25MG), geranylgeranyl pyrophosphate (Sigma-Aldrich, G6025), palmitic acid (Sigma-Aldrich, P0500), palmityl-carnitine (Sigma-Aldrich, P1645). If not stated in the legend, concentrations were 500 µM ligand, 100 µM MiD49, and 50 µM MiD51.

MiD49 or MiD51 was incubated with ligand for 1hr at 37 °C for 1 hr in S200 buffer, then mixed with the Native PAGE Sample buffer (Thermo Fisher Scientific, catalog BN2003). The samples were separated on a NativePAGE Novex 3%–12% Bis-Tris protein gel system (Thermo Fisher Scientific, catalog BN1003BOX) according to the manufacturer’s instructions. In brief, the electrophoresis was performed with 1× NativePAGE Running Buffer (Thermo Fisher Scientific, BN2001) and blue cathode buffer (containing 0.002% G-250). BN PAGE gel was destained in distilled water overnight, and band intensity was analyzed using ImageJ software.

### GTPase assay

DRP1 (0.75 µM) was mixed with indicated concentrations of MiD49, 51, MFF and/or actin filaments in S200 buffer. Sample were incubated at 37 °C for 5 min. At this point, GTP was added to a final concentration of 500 µM to start the reaction at 37 °C. Reactions were quenched at designated time points by mixing 15 µL sample with 5 µL of 125 mM EDTA in a clear, flat-bottomed, 96-well plate (Greiner, Monroe, NC). Six time points were acquired for all conditions, either in a 12 min time range, or in a 45 min time range depending on reaction speed. Released phosphate was determined by addition of 150 µL of malachite green solution as previously described ^26^ Absorbance at 650 nm was measured 15 min after malachite green solution incubation. GTP hydrolysis rates were determined by plotting phosphate concentration as a function of time in the linear phase of the reaction.

### Velocity Analytical Ultracentrifugation

Analytical ultracentrifugation was conducted using a Beckman Proteomelab XL-A and an AN-60 rotor. For sedimentation velocity analytical ultracentrifugation, MiD49/51 (5µM) in S200 buffer (65 mM KCl, 1 mM MgCl2, 0.5 mM EGTA, 1 mM DTT, 20 mM HEPES, pH 7.4) was centrifuged at either 5,000 (for oligomer) or 35,000 (for monomer) rpm with monitoring at 280 nm. Data analyzed by Sedfit to determine sedimentation coefficient, frictional ratio, and apparent mass. Sedimentation coefficient reported is that of the major peak (at least 80% of the total analyzed mass) at OD_280_.

### MANT-ADP assay

MANT-ADP assay was performed as previously described with modifications^20^. Fluorescence measurements of MANT-ADP (Sigma-Aldrich, 19511) were performed using a 96-well fluorescence plate reader (Infinite M1000; Tecan, Mannedorf, Switzerland) at room temperature in S200 buffer (65 mM KCl, 1 mM MgCl2, 0.5 mM EGTA, 1 mM DTT, 20 mM HEPES, pH 7.4). Samples were excited at 355 nm, and fluorescence emission was monitored at 448 nm. For titrations, MANT-nucleotide was held constant at 300 nM and the protein concentration was varied as indicated, measurements were conducted after 30 minutes incubation at room temperature.

### High-speed pelleting assay

Interactions between DRP1, actin and MiD49 were tested in the S200 buffer; 1.3 µM DRP1, 1 µM actin, and 4 µM MFF were mixed as described and were incubated for 1 hr at room temperature in a 100 µl volume. After incubation, samples were centrifuged at 80,000 rpm for 20 min at 4°C in a TLA-100.1 rotor (Beckman). The supernatant was carefully removed. Pellets were washed three times with S200 buffer and then resuspended in 100 µl of SDS–PAGE sample buffer and resolved by SDS–PAGE (LC6025; Invitrogen, Carlsbad, CA). Gels were stained with Coomassie Brilliant Blue R-250 staining (1610400, Bio-Rad, Hercules, CA), and band intensity was analyzed using ImageJ software.

### HPLC analysis of long-chain Acyl-CoA esters

HPLC analysis was performed as previously described with modifications^44^. A Betasil C18 column (150 x 4.6 mm) from Thermo (0711365H; Thermo Fisher Scientific, Waltham, MA) was used. The two mobile-phase solvents were 25 mM KH_2_PO_4_, pH 5.3 (pump A) and acetonitrile (pump B). Column was equilibrated in 95% pump A/5% pump B at 2 mL/min. A discontinuous gradient for elution of long-chain acyl-CoA esters was divided into four steps. Step 1: 95%/5% to 70%/30% over 5 min. Step 2: 70%/30% to 60%/40% over 2.5 min. Step 3: 60%/40% to 54%/46% over 4.5 min. Step 4: 54%/46% to 38%/62% over 2.5 min, then held for an additional 2.5 min at 38%/62% before re-equilibration by a 5 min reversed-flow gradient. The volume of the sample injected was 200 µL. The acyl-CoA esters were detected at. 260 nm. Quantitation was based on peak areas.

### Phosphate Assay

Fractions from Superose 6 size exclusion chromatography were assayed for phosphate content as follows. The fraction (0.4 mL) was mixed with 0.1 mL of 70% perchloric acid (Sigma-Aldrich 244252) in a 13×100 mm glass tube, and heated to 190°C for 20 min. After cooling, the following were added: 0.6 mL water, 0.25 mL molybdate solution (1.25% ammonium molybdate in 2.5 N sulfuric acid), and 0.06 mL Fiske-Subbarow reducer (Sigma-Aldrich 46345, 16% solution in water). Sample heated to 90°C for 20 min. After cooling, absorbance at 820 nm recorded. Phosphate content determined against standard curve of sodium phosphate, and converted to coenzyme A content by dividing by 3. This assay provides linear detection of inorganic phosphate from 1.5-40 nmole, and displays near-100% detection of known amounts of palmitoyl-CoA.

### Negative-stain transmission electron microscopy

Negative-stain TEM grids of purified MiD49 and MiD51 were prepared following an established protocol (Booth et al. PMID: 22215030). Briefly, 4ul of the sample at a concentration range of 0.1 – 0.4 mg/ml was applied to a glow-discharged 400 mesh copper grid coated with a continuous thin carbon film (prepared in house), blotted with filter paper, and then stained with freshly prepared 0.75% (w/v) uranyl formate. Grids were visualized at room temperature using a Tecnai T12 Spirit (FEI) equipped with an AMT 2k x 2k side-mounted CCD camera and operated at a voltage of 100 kV. Images were recorded at a nominal magnification range of 120,000-150,000x at the sample level with a calibrated pixel size range of 5.28-4.22 Å per pixel. Particle dimensions were measured by cropping 150×150 nm boxes from the original fields, and then determining the length of the longest and shortest axis for each particle manually.

### Cell culture and transfections

Human cervical cancer HeLa cells were purchased from ATCC (CCL-2) and grown in DMEM (Corning; 10-013-CV) supplemented with 10% fetal bovine serum (F4135; Sigma). Cells were cultivated at 37°C with 5% CO_2_. For plasmid transfections, cells were seeded at 4 × 10^5^ cells per 35 mm well 24 h before transfection. Transfections were performed in OPTI-MEM medium (Life Technologies; 31985062) with 2 µl of Lipofectamine 2000 (Invitrogen; 11668) per well for 6 h, followed by trypsinization and replating onto coverslips or glass-bottomed dishes (MatTek Corporation; P35G-1.5-14-C) at a cell density of ∼2 × 10^5^ cells per well. For MiD51 plasmid transfections carried out in wildtype HeLa cells 50 ng of each plasmid were transfected, respectively. For live-cell time-lapse acquisition, 100 ng of mito-plum were co-transfected with 50 ng of the respective MiD49 or 51 constructs.

For siRNA transfections, 1 × 10^5^ cells for control knockdown and 1.5 x 10^5^ cells for MiD49/51 or DRP1 knockdown were plated on 6-well plates, and 2 µl RNAi max (Invitrogen; 13778) and 63 pmol of each siRNA was used per well. Since MiD51 constructs were not sequence-modified for RNAi-resistance, 150 ng of the respective GFP-tagged MiD51 plasmids were used in MiD49/51 siRNA-treated cells. For these rescue experiments, siRNA-treated cells were transfected with 150 ng of respective MiD51 plasmids and 2 µl Lipofectamine 2000 ∼48 h after siRNAs have been transfected. Overall, cells were fixed and analyzed 72 h post siRNA-transfection.

### Western blotting and antibodies

For preparation of whole-cell extracts, confluent cell layers in 35 mm dishes were washed 3× with phosphate-buffered saline (PBS), lysed using ∼350 µl of 1×DB (50 mM Tris-HCl, pH 6.8, 2 mM EDTA, 20%glycerol, 0.8% SDS, 0.02% bromophenol blue, 1000 mM NaCl, 4 M urea), and boiled for 5 min at 95°C, and genomic DNA was sheared using a 27×G needle. Proteins were separated by standard SDS-PAGE and transferred to a PVDF (polyvinylidine difluoride) membrane (Millipore). The membrane was blocked with TBS-T (20 mM Tris-HCl, pH 7.6, 136 mM NaCl, and 0.1% Tween-20) containing 3% bovine serum albumin for 1 h and then incubated with the primary antibody solution at 4°C overnight. Primary antibodies used were as follows: MiD51 (rabbit; 20164-1-AP; Proteintech; 1:1000), GFP (rabbit, self-made,1:1000), DRP1 (mouse; 611112; BD Transduction Laboratories; 1:1000), GAPDH (G-9, mouse; Santa Cruz Biotechnology; 1:500), actin (mouse; mab1501R; Millipore; 1:1000), tubulin (DM1-α, mouse; T9026; Sigma; 1:10,000). After being washed with TBS-T, the membrane was incubated with horseradish peroxidase (HRP)-conjugated secondary antibodies (goat anti mouse #1721011; Bio-Rad or goat anti rabbit #1706515; BioRad) for 1 h at room temperature. Chemiluminescence signals were detected upon incubation with ECL Prime Western Blotting Detection Reagent (45-002-40; Cytiva Amersham) and recorded with an ECL Chemocam imager (SYNGENE G:BOX Chemi XRQ). For LI-COR Western blots, membranes were incubated with either IRDye 680 goat anti-mouse (926-68070; LI-COR), IRDye 800CW goat anti-rabbit (926-32211; LI-COR), or IRDye 680 donkey anti-chicken (926-68075) secondary antibodies for 1 h at room temperature. Signals were detected using the LI-COR Odyssey CLx imaging system.

### Immunofluorescence staining

For immunolabeling of proteins of interest, cells were seeded subconfluently on fibronectin (F1141; Sigma)-coated (1:100 in PBS) coverslips (72222-01; Electron Microscopy Sciences) in 35 mm dishes and allowed to spread overnight. On the following day, cells were fixed in pre-warmed 4% paraformaldehyde (PFA; 15170; Electron Microscopy Sciences) in PBS for 20 min followed by three PBS washes. Then, cells were permeabilized with 0.1% Triton X-100 in PBS for 1 min and washed 3x with PBS. Before antibody staining, cells were blocked with 10% calf serum in PBS for ∼30 min. Primary antibodies were diluted in 1% calf serum in PBS and incubated for 1 h. Mitochondria were visualized using a primary antibody against the OMM protein Tom20 (rabbit; ab78547; Abcam; 1:200). DRP1 was stained using DRP1 primary antibody (mouse; 611112; BD Transduction Laboratories; 1:50). Coverslips were washed several times in PBS and incubated with secondary antibody solution for 45 min. Either anti-rabbit Texas Red (TI-1000; Vector Laboratories; 1:300), anti-mouse Texas Red (TI-2000; Vector Laboratories; 1:300) or anti-rabbit Alexa Fluor 405 (A31556; Invitrogen; 1:300) were used as secondary antibodies. Coverslips were washed in PBS and fixed on glass slides using ProLong Gold antifade mounting media (P36930; Invitrogen).

### 2-bromopalmitate treatment

A 100 mM stock of 2-bromopalmitic acid (2-BP, Sigma-Aldrich 248422) or methyl-palmitate (MP, Sigma-Aldrich P5177) was made in DMSO, and diluted to 150 µM in DMEM+10% fetal bovine serum that had been pre-equilibrated overnight to 37°C/5% CO_2_ immediately prior to use. HeLa cells (80-90% confluent) were treated for 1 hr with this 150 µM 2-BP stock or MP as control. Cells were then fixed and stained as indicated above.

### Confocal microscopy

Imaging was performed on a Dragonfly 302 spinning-disk confocal (Andor Technology) on a Nikon Ti-E base and equipped with an iXon Ultra 888 EMCCD camera, and a Zyla 4.2 Mpixel sCMOS camera, and a Tokai Hit stage-top incubator was used. A solid-state 405 smart diode 100-mW laser, solid-state 488 OPSL smart laser 50-mW laser, solid-state 560 OPSL smart laser 50-mW laser, and a solid-state 637 OPSL smart laser 140-mW laser were used (objective: 100 × 1.4 NA CFI Plan Apo; Nikon). Images were acquired using Fusion software (Andor Technology). Live-cell imaging was performed in DMEM (21063-029; Life Technologies) with 25 mM D-glucose, 4 mM D-glutamine, and 25 mM HEPES, supplemented with 10% FBS (F4135; Sigma) on glass-bottomed dishes (MatTek Corporation; P35G-1.5-14-C).

### Analysis of MiD51 localization patterns on mitochondria

HeLa WT cells were transfected with 50 ng of either GFP-tagged MiD51 WT or various MiD51 mutants harboring a single amino acid exchange within their putative acyl-CoA binding pocket. After ∼6 h of transfection, cells were plated on fibronectin-coated coverslips, fixed and stained for Tom20 on the following day. GFP-fusion expression level was assessed by the detectability of GFP signal upon short (<100 msec, <50% laser power) or long (500 msec, 100% laser power) exposure, binning cells into high expression and low expression categories, respectively. Cells were classified in a blinded manner. For low expression, classification was for GFP pattern on mitochondria (punctate or non-punctate). For high expression, classification was for mitochondrial morphology (elongated, collapsed). Respective fractions of cells [%] were plotted in a stacked bar graph. Results combine three independently performed replicates of this experiment.

To assess the different localization patterns of MiD51 constructs in a more quantitative manner, the % coverage of the mitochondrial surface by GFP signal, as well as the average size of the GFP signal on the mitochondrion, were determined using ImageJ. 20 µm^2^-sized ROIs with representative MiD51 localization were auto-thresholded for the Tom20 staining and converted to binary masks. GFP signals were first processed by background subtraction using ImageJ (math → subtract → value: 500), then converted to a binary mask. Binary images were analyzed using the “analyze particles” tool with settings as follows: size (pixel2) 0.05–infinity, circularity 0.00–1.00. MiD51: Mito area ratios [%] as well as MiD51 average sizes [µm^2^] were plotted as box-and-whiskers plots using Microsoft Excel. Data corresponds to three independent experiments.

### Mitochondrial area quantification

Cells were silenced for both MiD49 and MiD51 and compared to cells treated with a scrambled siRNA. 48 h after siRNA transfections, cells were transfected with 150 ng of GFP-tagged MiD51 construct. Transfected cells were re-plated onto fibronectin-coated coverslips, fixed and stained for Tom20 on the following day. To avoid adverse effects of high MiD51 expression on mitochondria morphology, only cells with low expression levels were used for mitochondrial area analysis (requiring 500 msec exposure with 100% power of the 488 nm laser for GFP detection). For mitochondrial area quantifications, a 20 µm^2^-sized ROI of resolvable mitochondria (Tom20 signal) was selected, auto-thresholded, converted to a binary mask and analyzed using the “analyze particles” plug-in in ImageJ to obtain the number of mitochondrial fragments and the area of each fragment per ROI. The data shown as mean mitochondrial area in µm^2^ was plotted in a bar graph combining the results of four biological replicates.

HeLa cells, treated with siRNAs for either DRP1, MiD49/51 or containing a scrambled sequence, were plated on fibronectin-coated coverslips, and treated with growth medium containing either 150 µM 2-BP or MP for 1 h. Afterwards, cells were washed, fixed, stained and subjected to mitochondrial area measurements essentially as described above.

### Quantification of mitochondrially-associated DRP1 puncta

HeLa cells knocked down for MiD49/51 with or without re-expression of MiD51 constructs were analyzed for DRP1 recruitment to mitochondria as well as DRP1 puncta size. Cells were fixed and stained for endogenous DRP1 in combination with Tom20. 20 µm^2^-sized ROIs containing resolvable mitochondria in spread cell areas were thresholded using the same contrast settings for the Tom20 staining. DRP1 stainings were first processed by background subtraction using ImageJ (math → subtract → value: 1500), then converted to an 8-bit image. To determine DRP1 recruitment to mitochondria, DRP1 fluorescence signals overlapping with corresponding Tom20 staining were measured using the ImageJ “Colocalization” plug-in with the following parameters: ratio 40% (0–100%), threshold channel 1: 50 (0–255), threshold channel 2: 50 (0–255), display value (0–255): 255. Colocalized pixels were then converted to a binary mask and analyzed using the “analyze particles” tool with settings as follows: size (pixel2) 0.05–infinity, circularity 0.00–1.00. DRP1: Mito area ratios [%] were plotted as box-and-whiskers plots using Microsoft Excel. Furthermore, average DRP1 puncta size [µm^2^] was determined from these binary masks using the “analyze particles” plug-in. Three independent experiments were performed.

### Data processing and statistical analyses

For biochemical assays, numerical data were processed and assembled with Kaleidagraph (Synergy Software) and Photoshop CS4 (Adobe). Data analyses were carried out in ImageJ and Excel 2010 (Microsoft). Statistical comparisons were performed with GraphPad Prism 6.01 (Dotmatics) using unpaired t test. A probability of error of 5% (p ≤ 0.05; * in figure panels) was considered to indicate statistical significance; **, ***, **** indicated p values ≤ 0.01, 0.001, and 0.0001, respectively.

For imaging data, brightness and contrast levels were adjusted using ImageJ software. Figures were further processed and assembled with Photoshop CS4. Data analyses were carried out in ImageJ and Microsoft Excel. All statistical analyses and p-value determinations were done using GraphPad Prism 6.01. Data sets were either compared using an unpaired Student’s t-test or a One-way Anova multiple comparisons (Dunnett’s) test, as indicated in respective figures. A probability of error of 5% (p ≤ 0.05; * in figure panels) was considered to indicate statistical significance. **, ***, and **** indicate p values ≤ 0.01, ≤0.001, and ≤0.0001, respectively.

### Data Availability

No datasets were generated in this study. Primary data that was analyzed in this study are included in this published article (and its supplementary information files), or available from the corresponding author on reasonable request. The latter data include the primary analytical ultracentrifugation files.

## Acknowledgements

We thank Asan Abdulkareem, Mac Pholo Aguirre Huamani, Jose Delgado, Sam Liu, Zdenek Svyndrych and Andreia Isabel Ferreira Verissimo for much appreciated help during these investigations. We also thank Ajit Divakaruni and Anthony Jones for advice on acyl-CoA extraction methods from cells, and Cy Ocala for stimulating interaction. This work was supported by NIH R35 GM122545 and R01 DK088826 to HNH, NIH P20 GM113132 to the Dartmouth BioMT, and by the Deutsche Forschungsgemeinschaft (DFG) fellowship KA5106/1-1 (418076373) to FK.

## Notes

### Competing Interest Statement

The authors have declared no competing interest.

