## Supplementary Figures for "Long-chain fatty acyl-coenzyme A activates the mitochondrial fission factors MiD49 and MiD51 by inducing their oligomerization"

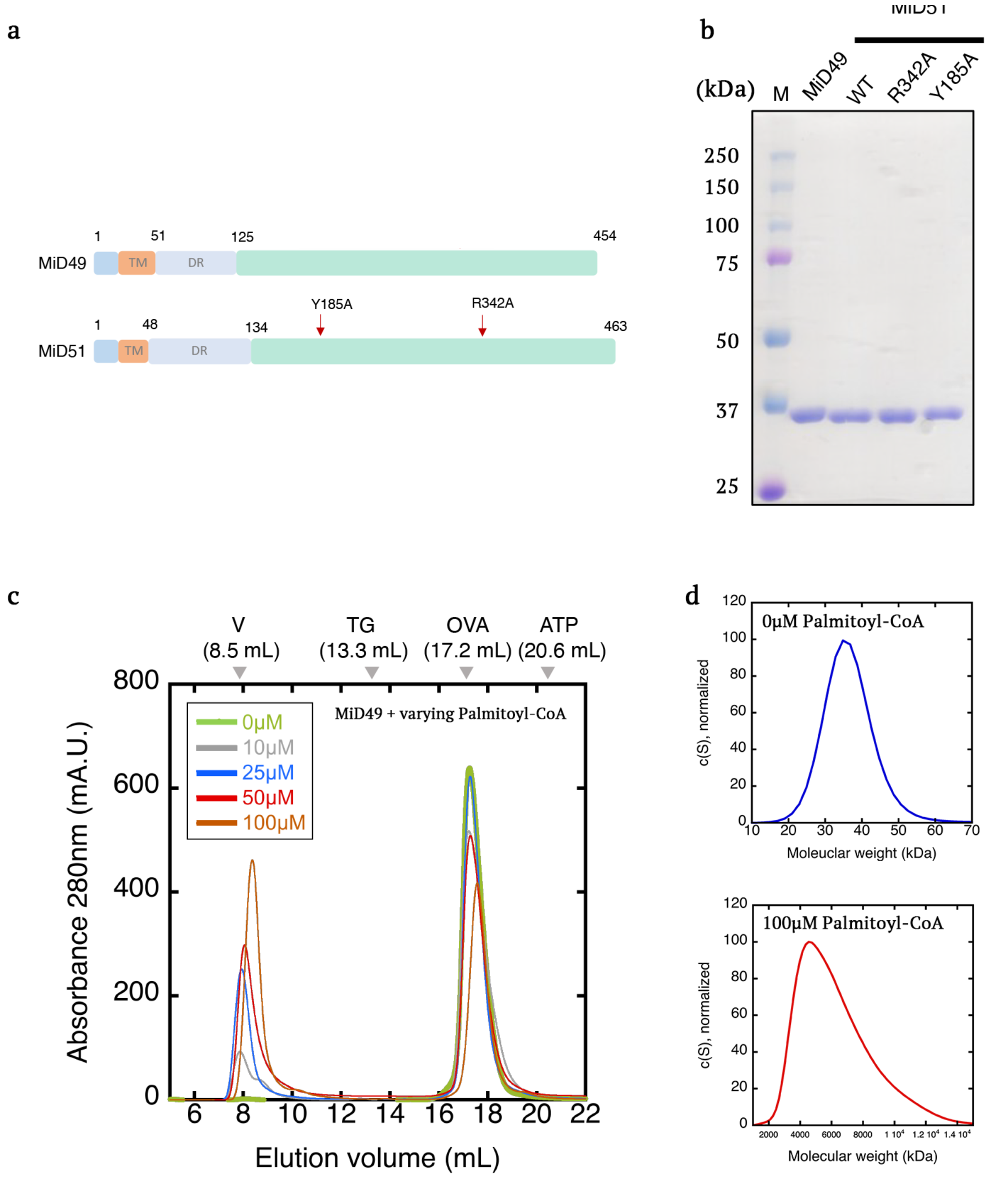


**Extended Data Fig. 1: Proteins used in this study, and effect of palmitoyl-CoA on MiD49 oligomerization.**
**a,** Bar diagrams of MiD49 and MiD51, showing the regions used in our biochemical studies: 125-454 for murine MiD49 (38.7 kDa), and 134-463 for human MiD51 (38.3 kDa). TM = transmembrane sequence, DR = disordered region. Green region = segment used for biochemical studies here, and for crystallization studies by others. **b,** Coomassie-stained SDS-PAGE of proteins used in this study. 2 μg protein loaded. Mass markers in kDa shown. **c,** Superose 6 size exclusion chromatography of MiD49 cytoplasmic region (100 μM) mixed with varying concentrations of palmitoyl-CoA. **d,** Velocity analytical ultracentrifugation of peak fractions from Superose 6 size exclusion chromatography of MiD49 cytoplasmic region (100 μM) incubated without or with palmitoyl-CoA (100 μM). Molecular masses calculated by these vAUC data: 36.4 kDa (without palmitoyl-CoA), and 4815 kDa (with palmitoyl-CoA).

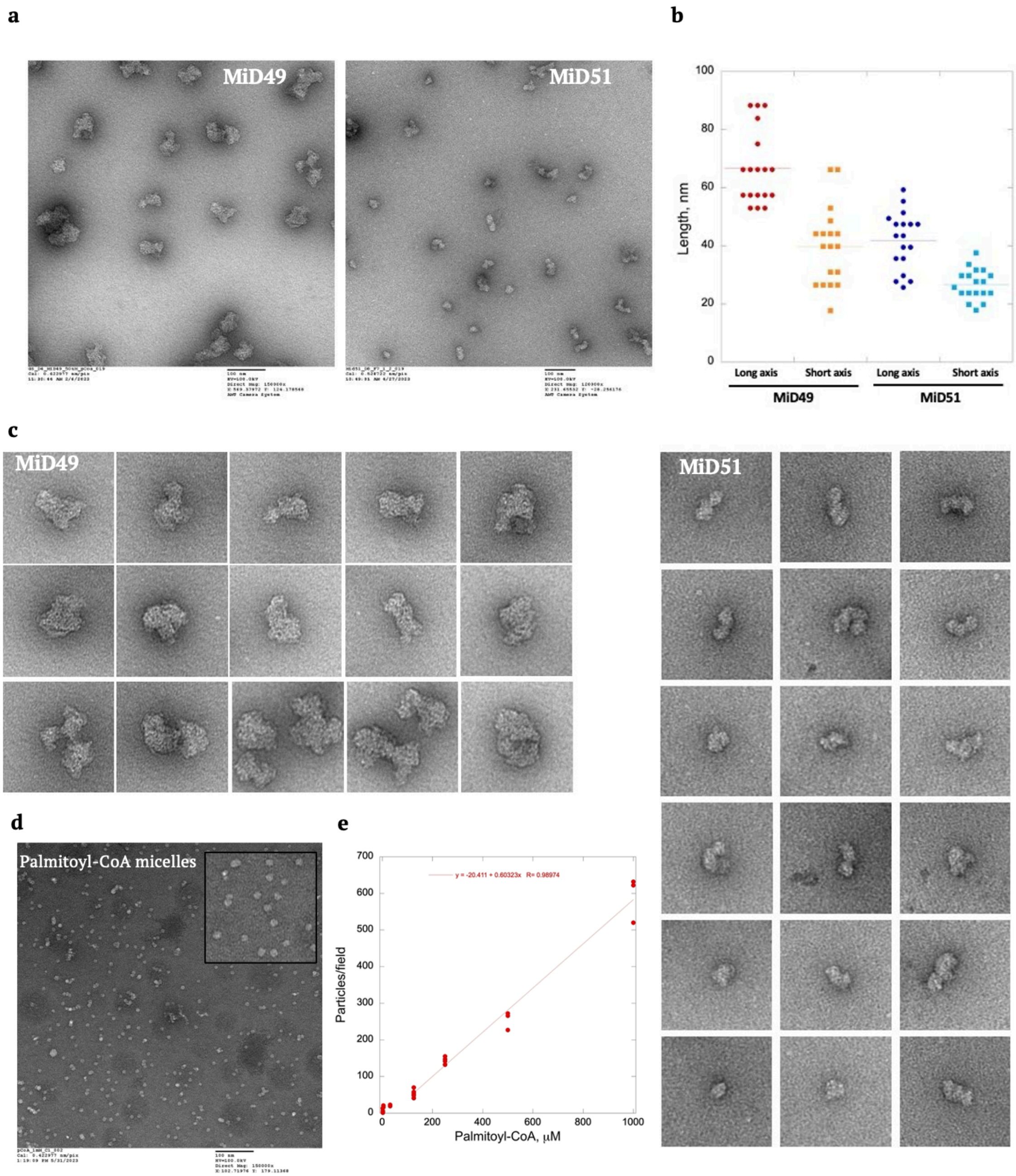


**Extended Data Fig. 2: Negative stain EM of MiD49 and MiD51 oligomers.**
Peak oligomer fractions from Superose 6 chromatography after mixing MiD49 (100 μM) or MiD51 (50 μM) with palmitoyl-CoA (50 or 500 μM, respectively).
**a,** Full-field examples for MiD49 and MiD51. **b,** Montage of MiD49 (left) or MiD51 (right) oligomer particles. Each image represents a 150 x 150 nm box. **c,** Graph of long axis and short axis lengths for 18 particles of MiD49 or MiD51 oligomers. **d,** Palmitoyl-CoA alone (1 mM) in same buffer as MiDs (10 mM Hepes pH 7.4, 65 mM KCl, 1 mM MgCl2, 1 mM EGTA, 1 mM DTT). Inset represents zoom of a 150 x 150 nm region. **e,** Graph of particles/field (800 x 800 nm field) versus palmitoyl-CoA concentration. 3-5 fields quantified for each concentration. X intercept for linear fit (after accounting for background particles, 8.2/field) represents critical micelle concentration (47 μM)

a b
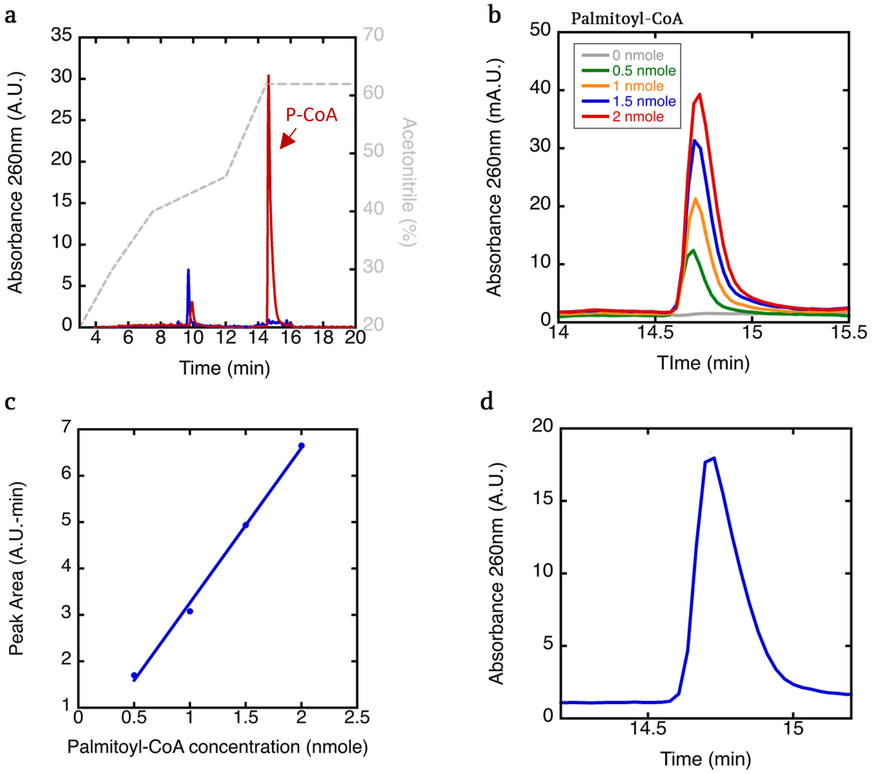


c d

e
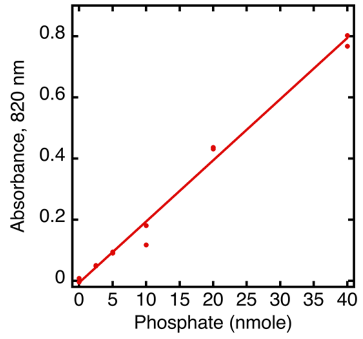


f

| | MiD49 | Phosphate | P-CoA | Ratio |
|---|---|---|---|---|
| Condition | µM | nmole | µM | P-CoA: MiD49 |
| 50 | 4.78 | 4.58 | 3.82 | 0.80 |
| 100 | 8.77 | 11.74 | 9.78 | 1.11 |
| 100 | 2.34 | 2.83 | 2.36 | 1.01 |

**Extended Data Fig. 3. Palmitoyl-CoA quantification in MiD49 oligomer fraction.**

**a,** Reversed-phase HPLC profile of 1.5 nmole palmitoyl-CoA (red) or solvent alone (blue). Position of palmitoyl-CoA (P-CoA) elution shown with arrow. Dashed gray line represents acetonitrile concentration. **b,** Palmitoyl-CoA peaks from four quantities of loaded palmitoyl-CoA (0, 0.5, 1, 1.5 and 2 nmole). **c,** Peak area as a function of palmitoyl-CoA loaded. **d,** Palmitoyl-CoA HPLC peak obtained when 200 mL of MiD49 oligomer fraction (Superose 6 oligomer fraction after incubating 100 µM MiD49 with 500 µM palmitoyl-CoA). Peak area, 3.42 A.U.-min, corresponding to 1.1 nmole P-CoA, or 5.5 µM. MiD49 concentration in this fraction (by Bradford assay), 5.3 µM. **e,** Standard curve of phosphate using the Fiske-Subbarow assay. **f,** Palmitoyl-CoA concentrations calculated from Fiske-Subbarow assay in which 400 µL oligomer fraction from Superose 6 was assayed. "Condition" refers to the µM amount of palmitoyl-CoA that was incubated with 100 µM MiD49 (37˚C, 1 hr) prior to separation of oligomers from monomers by Superose 6. MiD49 concentration determined by Bradford assay

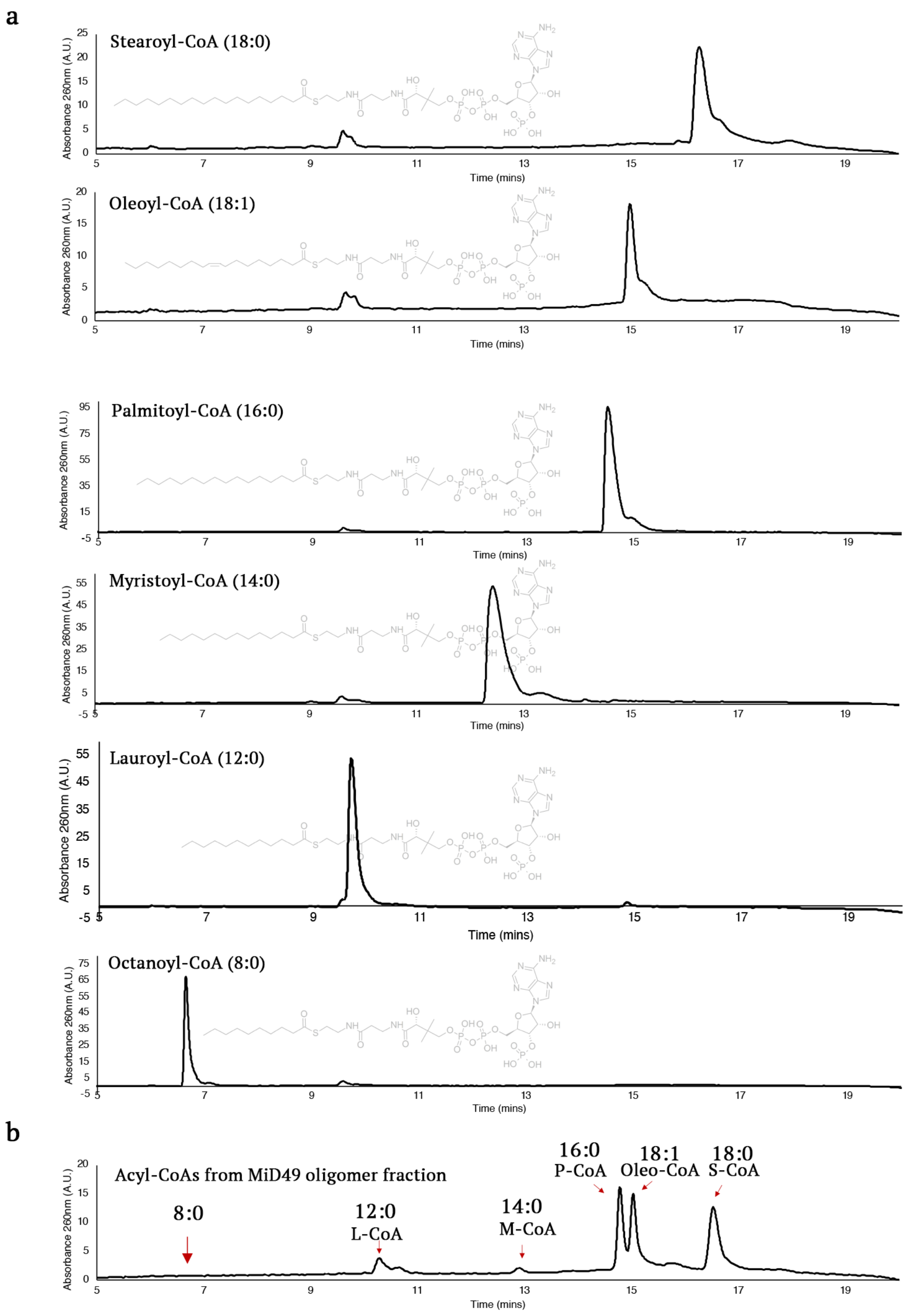


**Extended Data Fig. 4. HPLC analysis of acyl-CoAs bound to MiD49 oligomer.**

**a,** Individual HPLC traces of each of the acyl-CoAs (4 nmole loaded). **b,** HPLC trace of the MiD49 oligomer fraction from the following experiment. MiD49 cytoplasmic construct (100 µM) was mixed simultaneously with 83.3 µM of six acyl-CoAs (stearoyl, oleoyl, palmitoyl, myristoyl, lauroyl, octanoyl) for 1 hr at 37˚C, then MiD49 oligomer isolated by Superose 6 chromatography. Acyl-CoA distribution was then analyzed in the oligomer fraction by reversed-phase HPLC.

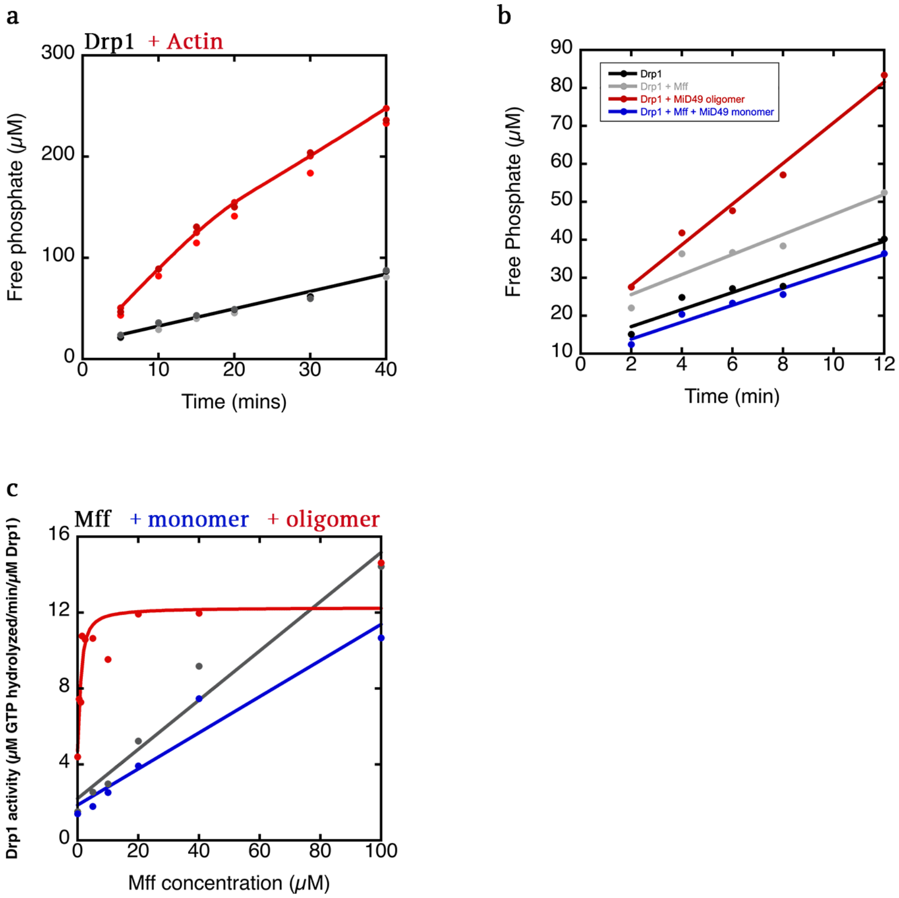


**<u>Extended Data Fig. 5. Effect of MiD49 and actin filaments on Drp1 activity</u>**

**a,** Drp1 GTPase assays (0.75 µM Drp1) alone (black points) or in the presence of 0.5 µM actin filaments (red). **b,** Drp1 GTPase assays (0.75 µM Drp1) alone (black points) or in the presence of 10 µM MFF alone (gray) or with 250 nM MiD49 oligomers (red) or monomers (blue) added. **c,** Effect of varying concentrations of MFF on Drp1 GTPase activity (0.75 µM Drp1) in the absence or presence of 250 nM MiD49 oligomers (red) or monomers (blue). Full curve to 100 µM MFF. Zoom to 20 µM MFF shown in Fig. 3f.

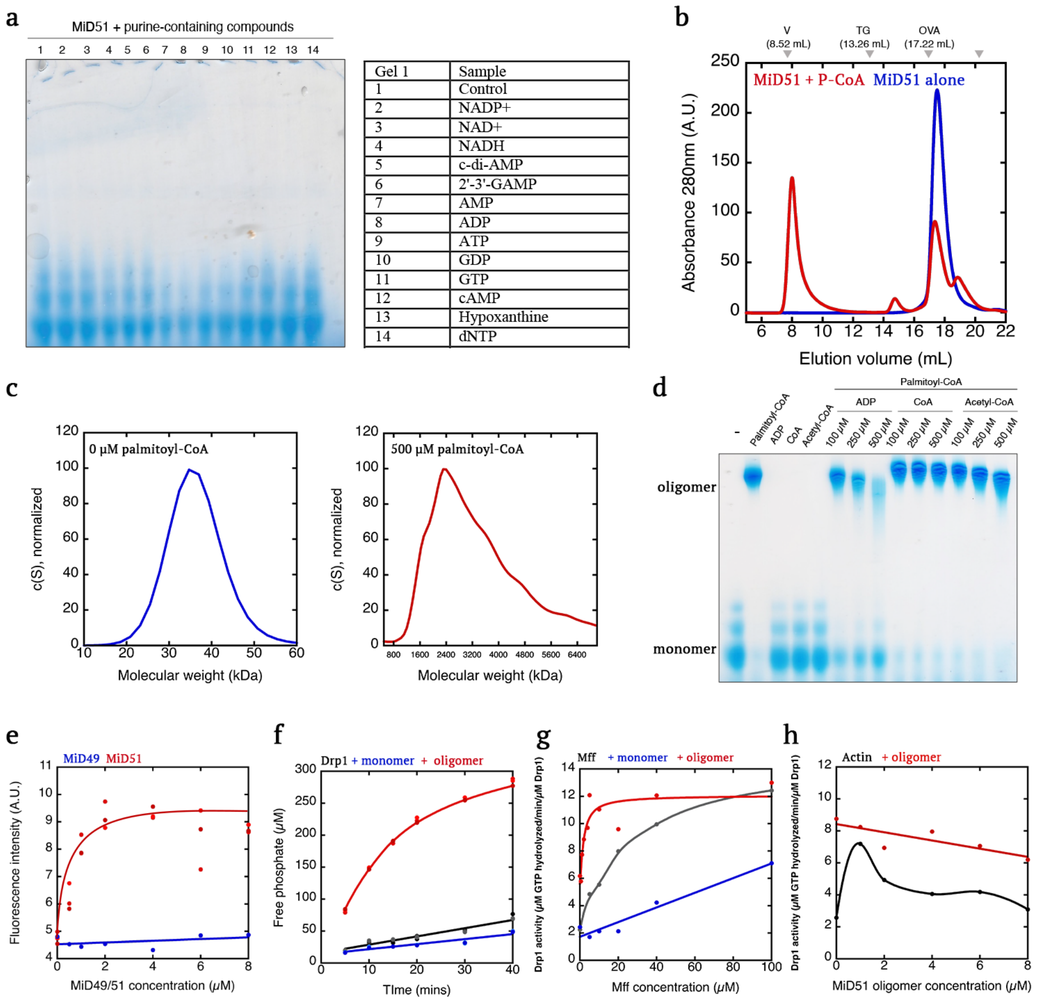


**Extended Data Fig. 6. Effects of LCACA on MiD51 biochemical properties**

**a,** Blue-native gel electrophoresis of MiD51 cytoplasmic region (50 µM) mixed with 500 µM of the indicated molecule. **b,** Size-exclusion chromatography (Superose 6) of MiD51 (50 µM) alone (blue) or in the presence of 100 µM palmitoyl-CoA (red). **c,** Velocity analytical ultracentrifugation of the MiD51 oligomer peak Fig. 4b (left) and the monomer MiD5 peak (right), showing calculated molecular masses.

**d,** Blue-native gel electrophoresis of MiD51 (50 µM) with 500 µM of the indicated molecules, or with 500 µM palmitoyl-CoA + the indicated concentrations of ADP, CoA or acetyl-CoA. **e,** MANT-ADP binding assay in which 300 nM MANT-ADP is mixed with the indicated concentrations of MiD51 or MiD49 and the fluorescence intensity monitored. **f,** Drp1 GTPase assays containing Drp1 alone (0.75 µM, black points) or in the presence of 0.5 µM MiD51 oligomers (red) or monomers (blue) added. **g,** Effect of varying concentrations of MFF on Drp1 GTPase activity (0.75 µM Drp1) in the absence or presence of 250 nM MiD51 oligomers (red) or monomers (blue). Full curve to 100 µM Mff. Zoom to 20 µM Mff shown in Fig. 5d. **h,** Effect of varying concentrations of actin filaments on Drp1 GTPase activity (0.75 µM Drp1) in the absence (black) or presence of MiD51 oligomers (blue).

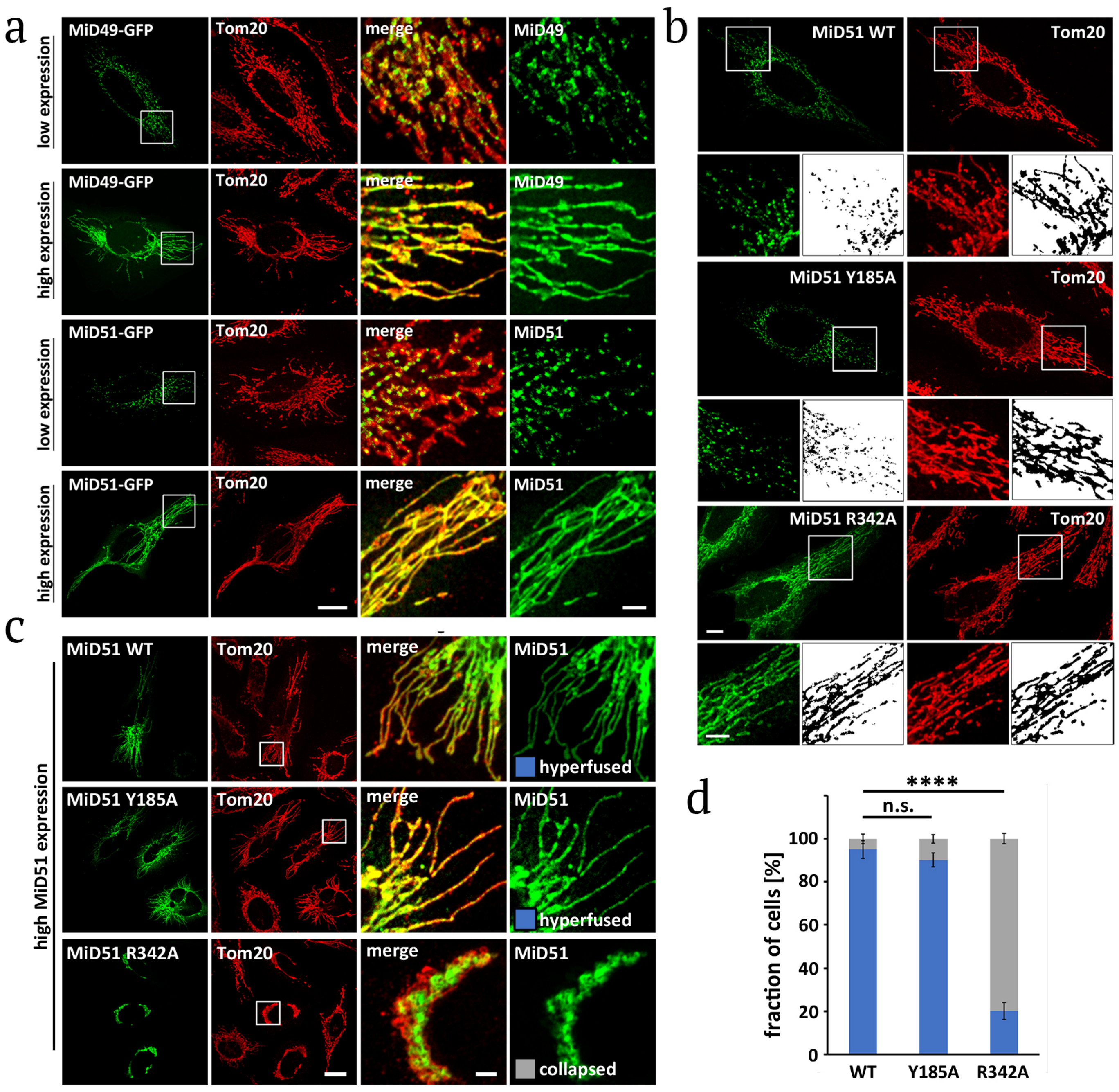
a
b
c
d
low expression
high expression
low expression
high expression
MiD49-GFP
Tom20
merge
MiD49
MiD51-GFP
MiD51
MiD51 WT
MiD51 Y185A
MiD51 R342A
high MiD51 expression
hyperfused
collapsed
****
n.s.
fraction of cells [%]
0
20
40
60
80
100
WT
Y185A
R342A

1

**Extended Data Fig. 7. Expression of MiD49 and MiD51 GFP fusion constructs in HeLa cells.** **a,** C-terminal GFP fusion constructs of MiD49 or MiD51 were transiently transfected into HeLa cells, then cells were fixed and stained for mitochondria (Tom20). At low expression levels (requiring 500 msec exposure at 100% laser power), the GFP fusions did not alter mitochondrial length, and displayed a punctate appearance on mitochondria. At high expression (100 msec exposure, 50% laser power), the GFP pattern on mitochondria was uniform and mitochondria were hyperfused. Scale bars: 20 mm (whole cell) and 3 mm (insets). **b,** Examples of image processing for quantification of GFP-MiD51 puncta size, and % mitochondrial area covered by GFP-MiD51. Boxed regions are shown in higher magnification below overview images. Binary masks of the respective inset images are shown on the right. These binary masks are representative of the images that were used for analyses in E and F. Scale bars are 10 mm in full images and 3 mm in insets. **c,** C-terminal GFP fusion constructs of the indicated MiD51 constructs were transiently transfected into HeLa cells, then cells were fixed and stained for mitochondria (Tom20). Cells were selected for high expression under the criteria described above. Images at right represent indicated boxed regions. Examples for WT and Y185 constructs represent hyper-fused phenotype quantified in panel D, while example for R342A mutant represents collapsed phenotype. Scale bars, 20 mm in full images and 3 mm in insets. **d,** Graph quantifying % cells displaying hyperfused (blue) or collapsed (gray) mitochondria. Representative examples of each pattern are shown below. Scale bar is 10 mm. N = 122, 82, and 85 cells were analyzed for MiD51 WT, Y185A, R342A, respectively. *** denotes p value of ⩽ 0.001 by ANOVA (Dunnett's multiple comparisons) test. n.s. = not significant (p value > 0.05). Bars in d represent standard error of the mean.

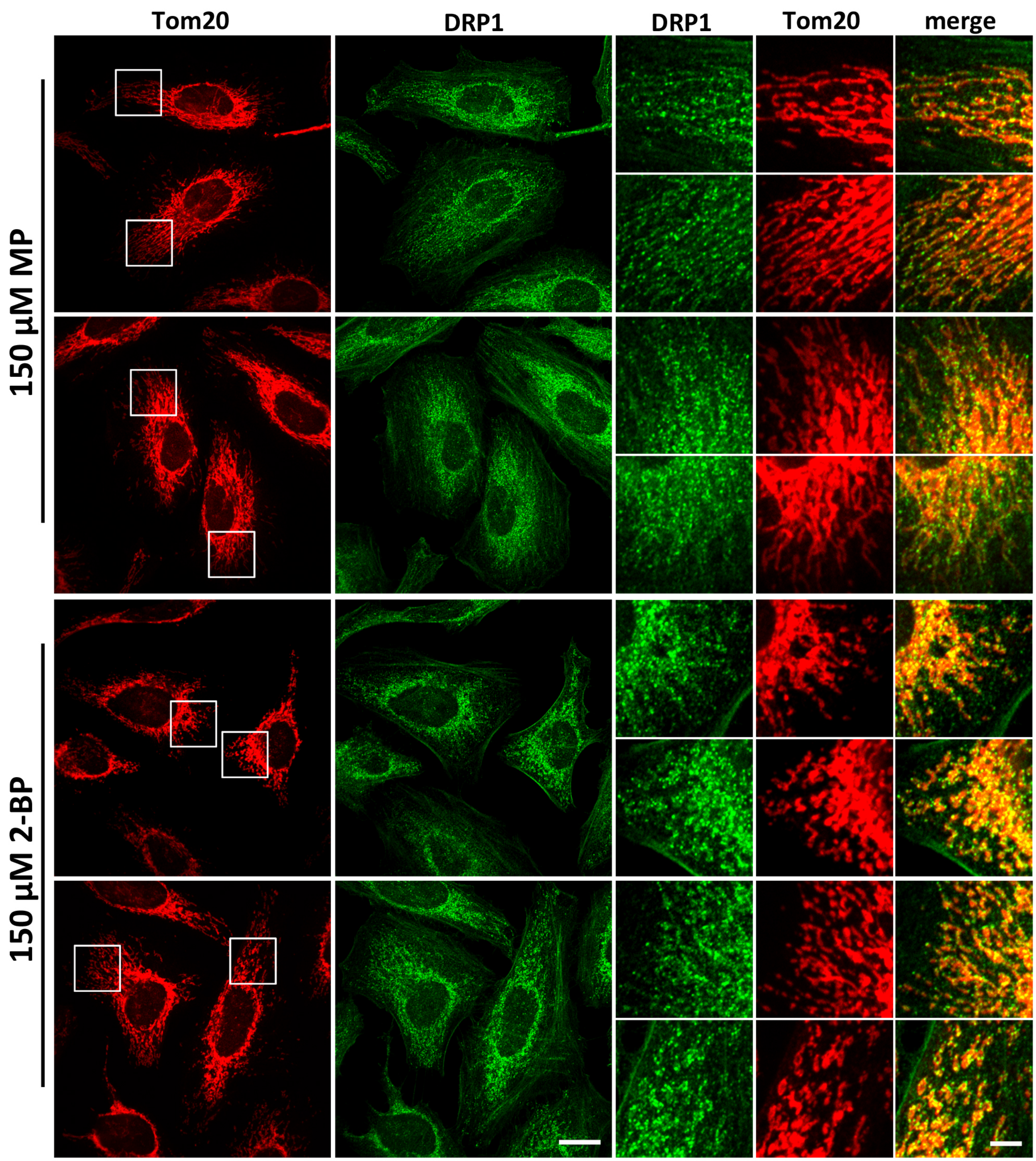


**Extended Data Fig. 8. Effect of 2-bromopalmitate on DRP1 mitochondrial puncta.**
HeLa cells were treated with either methyl-palmitate (MP) or 2-bromopalmitate (2BP) for 1-hr, then fixed and stained for mitochondria (Tom20, red) and DRP1 (green). Images at right represent zooms of indicated boxes at left. Two representative examples for either treatment are shown.

### Movies

**Movie 1. MiD49-GFP dynamics.** HeLa cell co-transfected with MiD49-GFP (green) and Mito-Plum (red). Scale bar in whole cell is 10 µm and 3 µm in inset.

**Movie 2. MiD51-WT GFP dynamics.** HeLa cell co-transfected with MiD51-GFP (green) and Mito-Plum (red). Scale bar in whole cell is 10 µm and 3 µm in inset.

**Movie 3. MiD51-Y185A GFP dynamics.** HeLa cell co-transfected with MiD51-Y185A-GFP (green) and Mito-Plum (red). Scale bar in whole cell is 10 µm and 3 µm in inset.

**Movie 4. MiD51-R342A GFP dynamics.** HeLa cell co-transfected with MiD51-R342A-GFP (green) and Mito-Plum (red). Scale bar in whole cell is 10 µm and 3 µm in inset.
